# A cortico-collicular pathway for motor planning in a memory-dependent perceptual decision task

**DOI:** 10.1101/709170

**Authors:** Chunyu A. Duan, Yuxin Pan, Guofen Ma, Taotao Zhou, Siyu Zhang, Ning-long Xu

## Abstract

Survival in a dynamic environment requires animals to plan future actions based on past sensory evidence. However, the neural circuit mechanism underlying this crucial brain function, referred to as motor planning, remains unclear. Here, we employ projection-specific imaging and perturbation methods to investigate the direct pathway linking two key nodes in the motor planning network, the secondary motor cortex (M2) and the midbrain superior colliculus (SC), in mice performing a memory-dependent perceptual decision task. We find dynamic coding of choice information in SC-projecting M2 neurons during motor planning and execution, and disruption of this information by inhibiting M2 terminals in SC selectively impaired decision maintenance. Furthermore, cell-type-specific optogenetic circuit mapping shows that M2 terminals modulate both excitatory and inhibitory SC neurons with balanced synaptic strength. Together, our results reveal the dynamic recruitment of the premotor-collicular pathway as a circuit mechanism for motor planning.

## INTRODUCTION

To survive in a dynamic environment with volatile sensory cues, it is often crucial for animals to maintain choice-related information in the gap between sensation and action. How the brain bridges past events with future actions is a fundamental question in neuroscience, and has been studied in animals using delayed-response motor planning tasks^1,2^. In these tasks, animals plan a response based on transiently presented sensory cues, but then need to withhold action and maintain this motor plan during a delay period without sensory cues. Using this classic paradigm, neural activity in the premotor cortex are shown to be critical for planning delayed responses in macaques^3,4^, rats^2,5,6^, and mice^7–10^.

Premotor cortex forms complex network with many cortical and subcortical areas. Recent work in head-fixed mice has started to reveal distinct contributions of premotor subpopulations based on their projection targets, such as the thalamus^11^, brainstem^7,12^, and cerebellum^13^. Thalamus-projecting premotor neurons contribute to the maintenance of persistent delay activity^11,12^; whereas brainstem-projecting neurons are more involved in motor execution^12^. Another major subcortical target of the premotor cortex is the midbrain superior colliculus (SC). Non-overlapping thalamus-projecting and brainstem-projecting premotor neurons both collateralize and project to the SC deep layers^12^. The SC also receives inputs from many other sensory and motor regions, and sends ascending and descending projections to various subcortical areas^14–16^, serving as a potential hub region that links sensation and action. Indeed, the SC has been implicated in planning and executing orienting responses in macaques^17–20^ and rats^5,21–23^. Given the vastly complex network associated with the premotor cortex and SC, it is crucial to specifically investigate the information routing in the core pathway linking these two key nodes in order to understand the circuit logic of motor planning. Although the cooperation between premotor cortex and SC was studied previously by inactivating the two regions during a delayed orienting task^5^, the approach likely also perturbed other parts of the network connected with the two regions. The information routing in the specific premotor-collicular pathway and its precise contribution to motor planning is yet to be investigated.

Here, we developed a delayed response task in head-fixed mice parametrizing the memory demands by varying the stimulus difficulty and delay duration, and employed projection-specific methods to examine the direct pathway from the secondary motor cortex (M2) to SC during memory-guided directional licking. Using multi-site optogenetic perturbations, we found concurrent involvement of M2 and SC in planning licking responses. To examine the temporal dynamics of information routing in the M2-SC pathway, we conducted *in vivo* two-photon calcium imaging from SC-projecting M2 neurons, and found that M2 sends progressively stronger choice-related information to SC, and the amount of choice signals accumulated in the M2-SC pathway correlates with animals’ response time. Furthermore, we used chemogenetic manipulation to specifically disrupt the M2-SC information flow by inhibiting M2 axonal terminals in SC, and found preferential impairment of mice’s performance in more demanding conditions with difficult stimuli and long delay durations, supporting a critical role of the M2-SC pathway in decision maintenance. Finally, we used cell-type specific optogenetic circuit mapping to probe the pattern of synaptic connectivity between M2 and SC, and found that M2 neurons synaptically target both excitatory and inhibitory neurons in SC with comparable strength, suggesting that the information sent from M2 to SC is further processed by balanced excitation-inhibition in SC local circuits. Our results reveal that the dynamic coding in the M2-SC pathway plays a causal role for decision-related processed in motor planning, identifying a core component of the cortico-subcortical network for memory-guided behavior.

## RESULTS

### M2-dependent memory-guided perceptual decision task

We trained head-fixed mice to perform a delayed-response auditory discrimination task, with varying perceptual difficulty and delay duration (Fig. 1). On each trial, mice were presented with auditory click trains of different rates (20-125 Hz). Click rates higher than 50 clicks/s indicate water reward from the left lick port; while lower lick rates indicate reward from the right port (Fig. 1a). Stimulus-response mapping was counterbalanced across animals. Mice need to discriminate the sensory cues, form and maintain a decision during a variable delay period, and respond by licking one of the two lick ports after an auditory “go” cue (Fig. 1b,c). Trials where animals lick before the “go” cue are considered violation trials and excluded from further analyses. Most previous studies using delayed-response tasks in head-fixed mice presented only two sensory stimuli to elicit two motor responses^8,10,24,25^, unable to vary stimulus difficulty. Here, we varied both perceptual difficulty (Fig. 1d) and delay duration (Fig. 1e,f) on a trial-by-trial basis, which allows us to examine the circuit mechanisms for motor planning with varying degree of memory demand. As delay duration increases, mice’s response time (RT) decreased from 567.1 ± 25.4 ms to 448.1 ± 26.8 ms (mean ± s.e.m. across animals; Fig. 1f), consistent with a motor preparation process^26^.

**Fig. 1.**
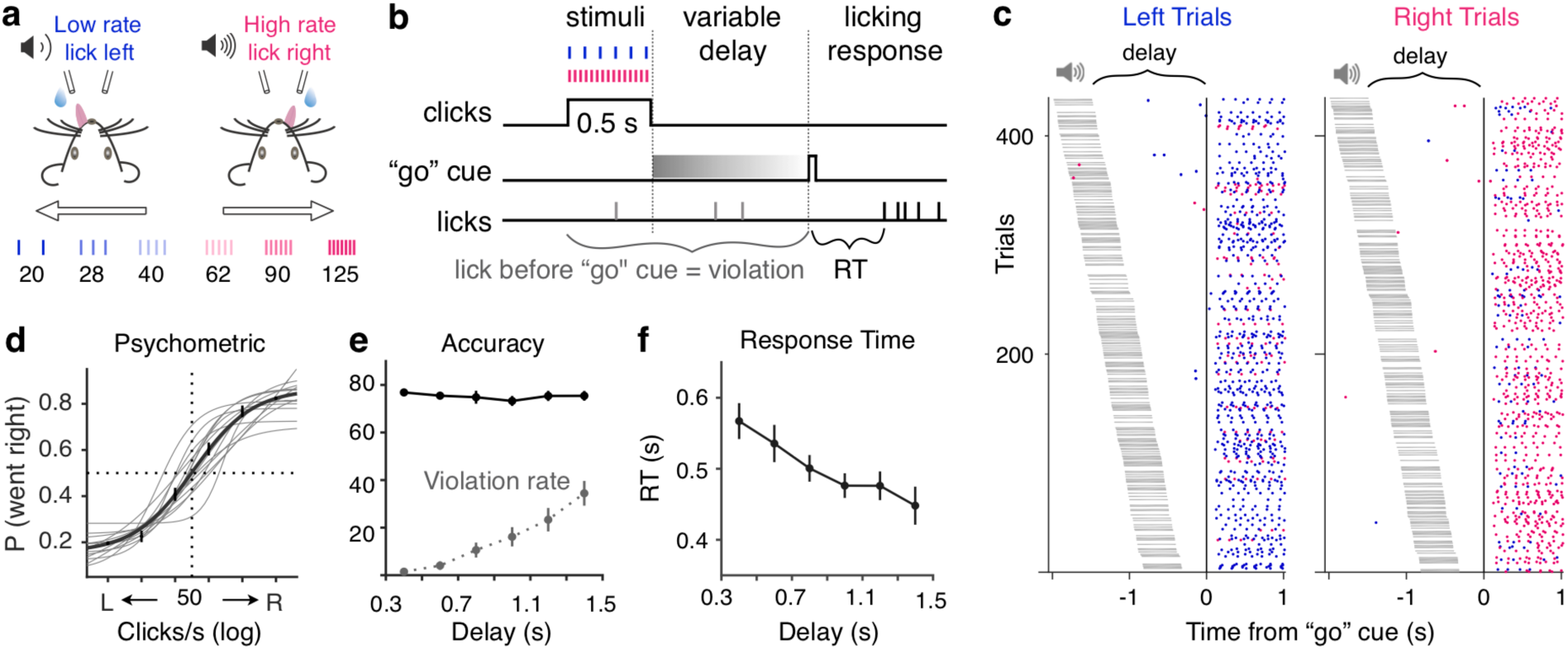
A memory-guided click rate discrimination task in head-fixed mice. **a**, Schematic of behavioral paradigm. **b**, Temporal structure of a task trial. A variable delay period sampled from a uniform distribution (0.3 - 1.5 s) is randomly assigned for each trial. Trials with premature licking are marked as violations, and excluded from further analyses. For non-violation trials, response time (RT) is defined as time between the onset of the “go” cue and animals’ first lick. **c**, Lick raster during an example session. Left licks (blue dots) and right licks (magenta dots) are plotted for left and right trials, aligned to the “go” cue, and sorted by delay durations. Timings of the sound are marked by gray horizontal lines. **d**, Sigmoid fit of psychometric performance. Thick line, mean fit; thin lines, individual mice (n = 14); error bars, s.e.m. across all trials. **e**, Accuracy (solid black) and violation rate (dotted gray) as a function of delay duration. Mean and s.e.m. across mice. **f**, Response time (RT) as a function of delay duration. Mean and s.e.m. across mice (median of RT distribution per mouse).

Converging evidence in head-fixed mice have implicated M2 as a cortical substrate for planning directional licking responses^7,8,10^. We thus tested whether our delayed-response auditory discrimination task also requires M2 activity using chemogenetic silencing (Supplementary Fig. 1). We expressed hM4Di, a designer receptor exclusively activated by designer drug (DREADD)^27^, in bilateral M2 neurons using a viral vector (AAV-Syn-hM4D(Gi)-mCherry), and conducted intraperitoneal (IP) injection of either Clozapine-N-oxide (CNO) or saline before each behavioral session (Supplementary Fig. 1a,b). The effect of inactivation was quantified by comparing task performance on CNO sessions to the corresponding saline control sessions (Methods). M2 inactivation impaired animals’ choice accuracy (*P*< 10^-3^, bootstrap, n = 15 sessions; Supplementary Fig. 1c,d), and increased animals’ RT on correct trials (RT increase = 128.1 ± 33.8 ms, *P*< 10^-3^; Supplementary Fig. 1e-g), with no difference between easy versus hard trials (*P*’s > 0.05). These results were not due to any non-specific effects of CNO IP injection alone (Supplementary Fig. 1i-l), and suggest that our task is M2-dependent.

### Involvement of SC in memory-guided directional licking

Although the midbrain SC has been extensively studied in the context of orienting behavior^14,15,28,29^, studies of SC’s functions for choice effectors beyond orienting are relatively rare^30,31^, and the precise contribution of the M2-SC pathway in motor planning remains unclear. To identify the anatomical M2-SC pathway potentially involved in memory-guided directional licking, we conducted anterograde tracing from M2 projection neurons to label the subregion of SC directly downstream of M2. We found dense axon terminals from M2 in the anterior lateral part of SC (Fig. 2a), a subregion of SC previously implicated in voluntary drinking^31^. Past work showed that M2 neurons that directly project to the brainstem reticular nuclei are involved in controlling directional licking^7^. Here, we labeled SC somas that are downstream of M2 using anterograde transsynaptic viral tracing^32^, and found that these SC neurons also project to brainstem reticular nuclei (Fig. 2b and Supplementary Fig. 2). These anatomical tracing experiments identify the lateral SC as a potential key node in the cortico-subcortical pathway controlling licking responses.

**Fig. 2.**
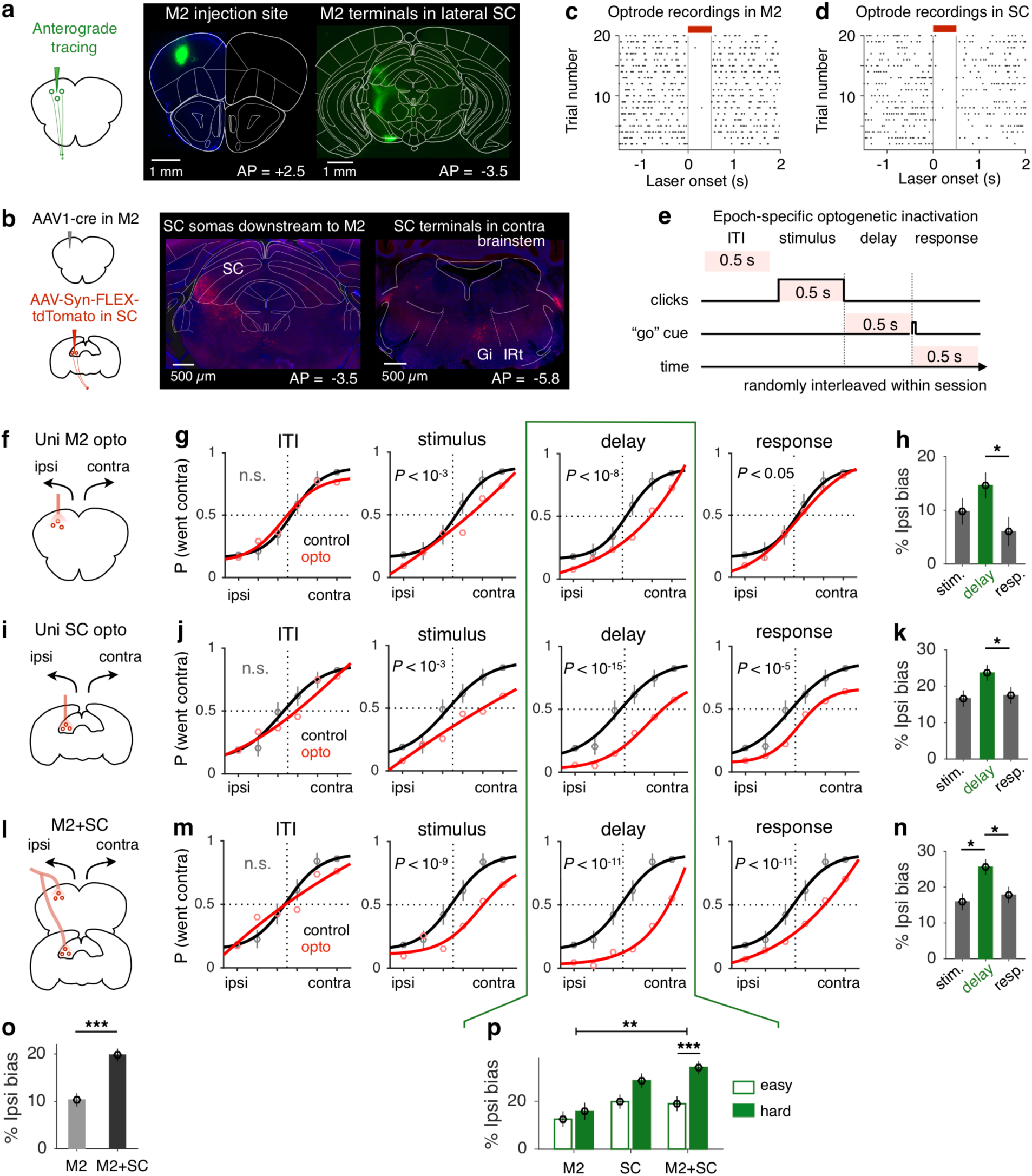
M2 and SC’s involvement in memory-guided directional licking. **a**, Anterograde tracing from M2 reveals projection terminals in lateral SC. **b**, SC somas downstream of M2 are labelled using transsynaptic virus (left). These SC neurons project to the contralateral brainstem (right). Gi, gigantocellular reticular nucleus. IRt, intermediate reticular formation. **c**, Acute extracellular recordings in awake mice to confirm optogenetic inactivation effect in M2. Spike activities in M2 neurons are aligned to laser onset over trials. Laser illumination period (0.5 s) is marked by the red bar. **d**, Similar to **c**, for SC. **e**, In each session, optogenetic inactivation during different behavioral epochs were randomly interleaved with no-laser trials (70%). The duration of laser stimulation was kept constant for all epochs (500 ms). **f**, Optogenetic inactivation of unilateral M2 (left or right). **g**, Data (circles, means and s.e.m. across 5219 trials concatenated across 12 sessions) and 4-parameter sigmoid fit (lines, for visualization only) for M2 inactivations during the inter-trial interval (ITI), stimulus, delay, or response period, compared to control trials. *P* values report ipsilateral bias after optogenetic inactivation, based on generalized linear mixed model fit (GLMM, logistic fit), lme4 package in R. **h**, Mean ipsilateral bias caused by M2 inactivation during different behavioral epochs; error bars, s.e.m. across trials. Between-epoch *P* values report corrected single-step multiple comparisons of GLMM results. \**P* < 0.05. **i-k**, Optogenetic inactivation of unilateral SC (5747 trials concatenated across 12 sessions), similar to **f-h**. **l-n**, Simultaneous optogenetic inactivation of M2 and SC on the same side (5425 trials concatenated across 12 sessions), similar to **f-h**. **o**, Mean ipsilateral bias after M2 inactivation or simultaneous inactivation of M2 and SC; error bars, s.e.m. across all trials and all epochs. Between-region *P* values report corrected single-step multiple comparison test of GLMM results. \*\*\**P* < 10^-3^. **p**, A difficulty-dependent impairment was observed only when M2 and SC neurons were both inactivated during the delay period (GLMM).

To test how M2 and SC are dynamically recruited for planning and executing decision-related licking responses, we separately inactivated M2 or SC neurons using optogenetics during different trial epochs, including the inter-trial interval (ITI), stimulus, delay, or response period (500 ms inactivation for all epochs; Fig. 2c-e and Supplementary Fig. 3). In each behavioral session, photostimulation was delivered unilaterally on 30% of randomly chosen trials to inhibit local activity (Fig. 2c,d; Methods). Performance on inactivation trials was compared to that on control trials in the same session. Different optogenetic conditions (ITI, stimulus, delay, or response period) were interleaved for all sessions to control for behavioral fluctuations across days. We used a generalized linear mixed model (GLMM) to quantify the effect of optogenetic inactivation on animals’ behavior while taking into account between-subject variance (Methods). We found that unilateral M2 inactivation led to a significant ipsilateral bias (contralateral impairment) when laser was delivered during the stimulus (ipsi bias = 9.9 ± 2.5%, *P* < 10^-3^, GLMM), delay (14.4 ± 2.4%, *P* < 10^-8^), or response period (5.8 ± 2.7%, *P* < 0.05), but not during the ITI ((−1.1 ± 3.3%, *P* = 0.62; Fig. 2f-h). The greatest impairment occurred when M2 was inactivated during the delay period (corrected single-step multiple comparison test of GLMM; Fig. 2h), suggesting that M2 plays a more important role during motor planning than during motor execution.

Because SC is classically associated with motor functions^28^, it is thus conceivable that SC may be more involved in motor execution than motor planning. On the contrary, similar to the M2 inactivation results, we found the greatest impairment of inhibiting SC activity following delay period inactivation (ipsi bias = 23.3 ± 2.1%, *P* < 10^-15^, GLMM), with significant effects also found after stimulus (16.0 ± 2.3%, *P* < 10^-3^) or response period inactivation (16.9 ± 2.3%, *P* < 10^-5^), but not during the ITI (4.2 ± 3.6%, *P* = 0.36; Fig. 2i-k). These results suggest that both M2 and SC are preferentially involved in motor planning in the context of directional licking responses.

To distinguish whether SC plays a permissive role relaying motor command from M2 to brainstem or a more active role, further processing cortical information to facilitate motor planning, we used a within-subject multi-site optogenetic perturbation approach to compare M2 inactivation with simultaneous inactivation of both M2 and SC during different behavioral epochs (Fig. 2l-p and Supplementary Fig. 4). Recordings using optrode in awake mice showed that the optimal laser stimulation parameters are different for M2 and SC neurons (Supplementary Fig. 3d,e; Methods). To ensure the temporal resolution of optogenetic silencing, we used the lower stimulation parameters for the simultaneous inactivation experiment, which resulted in complete inhibition of M2 activity and partial inhibition of SC activity. Therefore, the simultaneous inactivation can only be fairly compared to M2 inhibition alone.

We used a single GLMM to fit the optogenetic perturbation effects across 3 types of brain region inactivation and 4 task epochs (ITI inactivation as control), with click rates, region, and epoch as fixed effects and random effects across animals (Supplementary Fig. 4a,c, Model 1, red curves; Methods). Single-step multiple comparisons of GLMM results revealed that simultaneous inactivation of M2 and SC resulted in a significantly greater bias than M2 inhibition alone (*P* < 10^-3^; Fig. 2o and Supplementary Fig. 4b,c), suggesting that SC makes additional contributions to planning directional licking responses beyond relaying commands from M2. Moreover, this analysis confirmed that M2 and SC are preferentially involved during the delay period, compared to the stimulus (*P* < 0.01) or the response period (*P* < 10^-3^, GLMM, Supplementary Fig. 4c). Finally, we tested whether optogenetic perturbation of the premotor-collicular circuit affected difficult trials more compared to easy trials by adding a three-way interaction term (region × epoch × difficulty) to the GLMM (Supplementary Fig. 4a,c, Model 2, blue curves). Including this interaction term significantly improved the overall model fit (*P* < 0.05, likelihood ratio test, Chisq). The difficulty effect was significant only when M2 and SC neurons were both silenced during the delay period (*P* < 10^-3^; Fig. 2p and Supplementary Fig. 4c), but not in any other condition (*P*’s > 0.05, overlapping red and blue curves in Supplementary Fig. 4c), consistent with the possibility that M2 and SC make concurrent contributions to decision maintenance.

### Projection-based imaging reveals choice-related information in the M2-SC pathway

To test if the projections from M2 to SC carry task-relevant information, we performed projection-specific *in vivo* two-photon calcium imaging from a subset of M2 neurons that project to the SC with retrograde expression of GCaMP6s (Fig. 3a; Methods)^33^. We imaged populations of apical dendritic trunks and somas of SC-projecting M2 neurons in randomly selected fields of views (627 ROIs, 9 sessions, 4 mice) during task performance, and found 64.0% of neurons to be side-selective (*P* < 0.01 for any behavioral epoch; Fig. 3b-c; Methods). Individual neurons show heterogeneous choice-related activity during pre-movement and movement periods (Fig. 3d and Supplementary Fig. 5a). To compare choice selectivity across different behavioral epochs for individual neurons, we conducted receiver operating characteristic (ROC) analysis^34^ on responses during the relevant time window for each behavioral epoch (Fig. 3e,f); and calculated the area under ROC curve (AUC) to indicate discrimination between contra-versus ipsi-lateral choices (Methods). At the population level, we compared distributions of AUC values and observed a progressively stronger contralateral preference as the trial unfolds, from the sound epoch, to the delay epoch, and finally the response epoch (*P* < 0.01 for each pair of epochs in the example animal, *P* < 10^-3^ for each pair of epochs for data across all animals, permutation tests; Fig. 3g and Supplementary Fig. 5b). This gradual increase of contralateral choice selectivity during the trial reflects an emerging motor command in the premotor-collicular pathway.

**Fig. 3.**
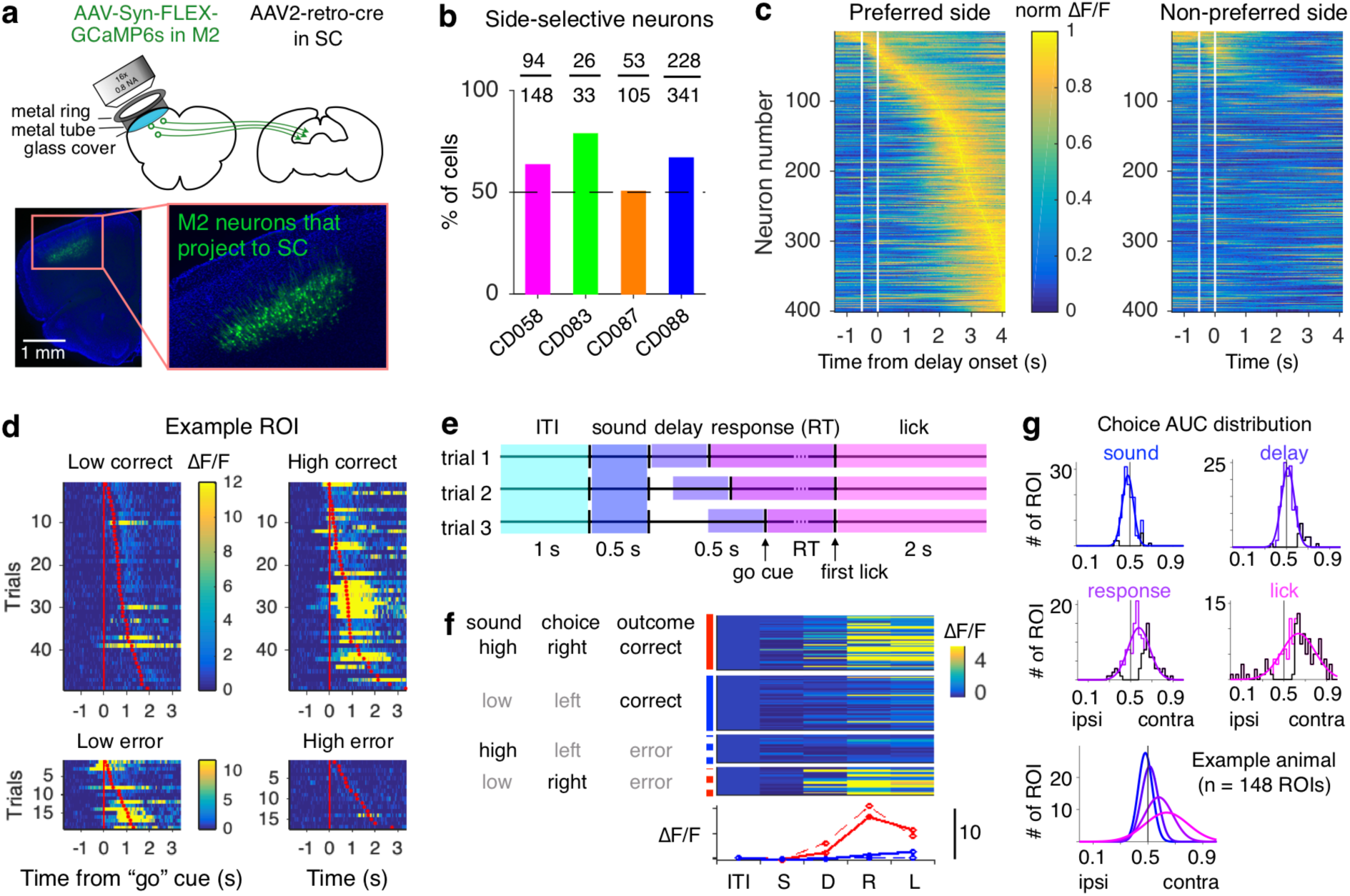
Projection-based imaging reveals choice-related information in the M2-SC pathway. **a**, Design for projection-based imaging experiments (top) and example histology that shows selective expression of GCaMP6s in SC-projecting M2 neurons (bottom). **b**, Percentage and actual number of side-selective neurons for different animals. **c**, Dynamics of all side-selective neurons separated by the choice of the animals, sorted by the time of peak activity on the preferred side. White lines mark the timing of the sound period. **d**, Responses (ΔF/F) of an example choice-selective ROI from one imaging session. Each row shows the response for one trial. Red ticks mark RT across trials. Each of the four panels shows one trial type. Diagonal panels correspond to the same type of choice. **e**, Schematic that shows the relevant time window of responses extracted for different behavioral epochs. **f**, Simplified response profile for the same ROI as **d**. Top, average activity across frames in each behavioral epoch for individual trials. Each block of trials correspond to one of the four trial types. Bottom, red and blue traces show the average ΔF/F responses for right and left trials (solid for correct, dashed for error trials) across behavioral epochs. **g**, Histograms and gaussian fits (smooth lines) to distributions of AUC values of all the ROIs from one example animal (n = 148 ROIs), during the sound, delay, response, and lick epochs. Black lines show individually significant neurons. Right, distributions across epochs are plotted together to reveal a gradual increase of contralateral selectivity over time.

The projection-based imaging method enabled us to sample a large number of neurons simultaneously with cell-type specificity (69.7 ± 23.4 ROIs/session), which in turn allowed us to decode animal’s choice on a single-trial basis from population activity in the M2-SC pathway. We used cross-validated linear classifiers to decode the amount of choice or sensory information during different behavioral epochs (Fig. 4a,b; Methods). Although the number of individually choice-selective neurons was small during the delay period (77/627, *P* < 0.05, ROC; Supplementary Fig. 5c), population-level delay activity contained significantly above-chance choice information in most sessions (6/9 sessions, *P* < 10^-3^; Fig. 4c). In addition, the amount of choice information was significantly higher than the amount of sensory information during the delay (bootstrap, *P* < 10^-3^). Consistent with single-neuron analyses, population decoding results also showed progressively stronger choice information over time. A repeated measure two-way ANOVA on decoding accuracy revealed a significant main effect of choice versus sensory information (*P* < 10^-3^), a significant main effect of task epoch (*P* < 10^-10^), and a significant effect of interaction (*P* < 10^-5^; Fig. 4c).

**Fig. 4.**
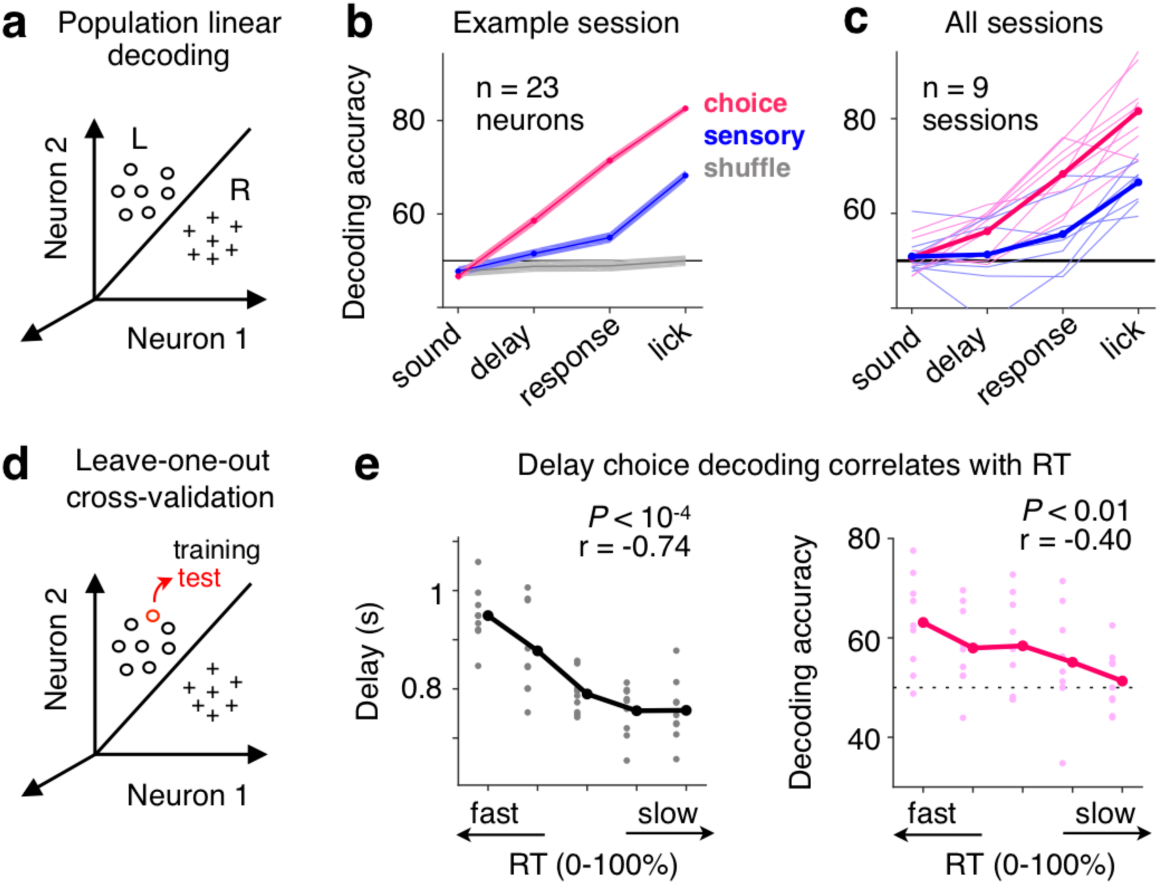
Linear classification of population activity predicts animal’s choice and RT. **a**, Schematic to illustrate the linear decoding method using population activity. Clouds of hypothetical population responses corresponding to different conditions (left or right choices) are linearly discriminated in high dimensional neural activity space. **b**, Classification accuracy of decoders trained on 23 simultaneously recorded neurons from one example session. Choice decoding corresponds to classification between left versus right choices; sensory decoding corresponds to classification between high versus low click rate trials; gray lines show performance on data with shuffled trial labels. Mean ± s.e.m. across 100 cross-validated samples. **c**, Decoding accuracy for all sessions (n = 9), similar to **b**. Thin lines, individual sessions; thick lines, mean across sessions. **d**, Schematic to illustrate leave-one-out cross-validation. **e**, Left, delay durations binned over each 20 percentile of response times (RT) across sessions (n = 9). Gray dots, individual sessions; black dots, mean across sessions. Right, delay activity choice decoding accuracy binned over each 20 percentile of RT. Light pink dots, individual sessions; Dark pink dots, mean across sessions.

To examine the relationship between the amount of accumulated choice information in the M2-SC pathway and animal’s behavior on a trial-by-trial basis (Fig. 4d), we evaluated the correlation between delay-period activity and animal’s response time (RT). We found that shorter RT tended to happen on trials with longer delay durations (r = −0.74, *P* < 10^-4^; Fig. 4e, left), consistent with a motor preparation process where movements are faster when more time is given to prepare for them^26^. Remarkably, the amount of choice information decoded from the SC-projecting M2 populations during the delay period was also negatively correlated with RT (r = −0.40, *P* < 0.01; Fig. 4e, right), suggesting that the strength of choice signals accumulated in the M2-SC pathway contributes to motor preparation. We fit a linear mixed effect model (LMM) to simultaneously quantify the effects of delay duration and delay neural activity in predicting RT (Methods), and found a significant effect of choice decoding accuracy (*P* < 0.05, LMM) beyond the predictive power of delay duration (*P* < 10^-3^). The effect of decoding accuracy in predicting RT remained the same after controlling for behavioral performance on each trial (Methods). Together, single-neuron and population analyses of projection-based two-photon imaging data reveal that M2 sends progressively stronger choice-related information to SC, and such information during the delay period predicts animals’ subsequent behavior.

### Causal contribution of the M2-SC pathway to decision maintenance

To test whether the choice signals in the M2-SC pathway play a causal role in behavior, we selectively inhibited the projections from M2 to SC using chemogenetics (Fig. 5a). We expressed hM4Di in bilateral M2 neurons and locally infused CNO or saline in bilateral SC via implanted cannulae (Supplementary Fig. 6; Methods). Different from a general error increase for all trials after M2 soma inactivation (Supplementary Fig. 1d), silencing M2 projections to SC preferentially impaired hard trials but not easy trials (*P* < 10^-3^; Fig. 5b,c). This effect is consistent with the difficulty-dependent effect following simultaneous optogenetic inactivation of both M2 and SC neurons during the delay period (Fig. 2p and Supplementary Fig. 4c), suggesting a specific role of the M2-SC pathway in decision-related processes. The variable delay period in our task allowed us to further assess the role of the M2-SC pathway on trials when animals needed to maintain decisions over a short versus long period of time. We found that the difficulty-dependent impairment only occurred on long-delay trials (*P* < 10^-3^) but not on short-delay trials (*P* = 0.61; Fig. 5d,e). A repeated-measures two-way ANOVA on error increase revealed a significant main effect of difficulty (*P* < 0.05); a significant interaction between difficulty and delay length (*P* < 0.05); and no effect of delay *per se* (*P* = 0.26). The delay- and difficulty-dependent impairment supports a critical role of the M2-SC pathway for decision maintenance.

**Fig. 5.**
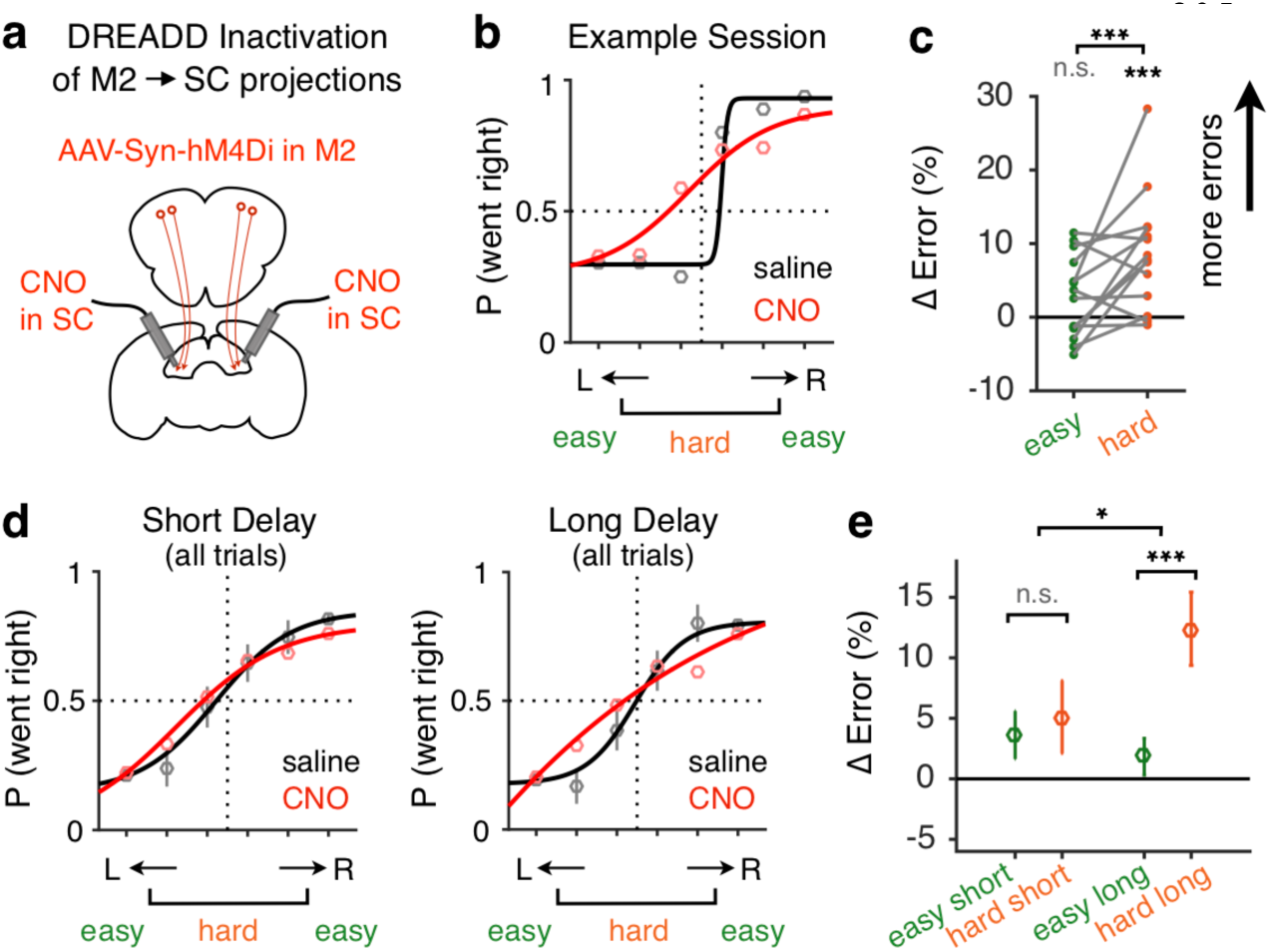
Inactivation of M2 terminals in SC impairs decision maintenance. **a**, Design for bilateral chemogenetic inactivation of M2 terminals in SC. **b**, Example SC infusion session (red) compared to saline control session (black) for one animal. Dots, data; lines, sigmoid fit. **c**, Difference in error rate between inactivation sessions (n = 14) and saline control sessions (n = 14) for easy (green) and hard (orange) trials. Gray lines connect data from the same session. **d-e**, Psychometric curves (**d**) and error rate increase (**e**) due to M2 terminal inhibition in SC, plotted separately for short delay (300 - 900 ms) and long delay trials (900 - 1500 ms). **e**, Mean and s.e.m. across 14 pairs of sessions. \**P* < 0.05; \*\*\**P* < 10^-3^; n.s., not significant; permutation test.

### M2 targets SC excitatory and inhibitory neurons with comparable synaptic strength

Our chemogenetic experiment selectively disrupted M2 axons in SC without perturbing other M2 projections. Therefore, the effects we observed here can be specifically attributed to the M2-SC pathway, and further processing of choice-related information in the SC may be essential for decision maintenance (Supplementary Fig. 4). We thus probed how this information may be transformed within the SC. Combining anterograde transsynaptic viral tracing and immunohistochemistry (Methods), we found that 48.3% of the SC neurons downstream of M2 are GABAergic (Fig. 6a), suggesting that M2 terminals target both excitatory and inhibitory neurons in the SC. To conduct cell-type-specific circuit mapping of M2-SC connections, we measured synaptic transmission in genetically labeled excitatory or inhibitory SC neurons using patch-clamp recording while optogenetically stimulating M2 terminals in brain slices (Fig. 6b,c; Methods). A train of 10 pulses of photostimulation was first delivered to test the responsiveness of SC neurons that receive M2 input (Fig. 6c). The percent of responsive neurons was similar in the excitatory (75%, 18/24) and inhibitory (63.6%, 28/44) SC populations (*P* > 0.05, chi-squared test; Fig. 6d, top left). For responsive neurons, we repeatedly delivered single-pulse stimulation to test the response latency, reliability (probability of response), and amplitude of excitatory postsynaptic potential (EPSP), and found no significant difference between the two populations (*P’s* > 0.05, permutation test; Fig. 6d). These data suggest that M2 neurons synaptically target both excitatory and inhibitory neurons in SC with comparable strength. The regulation of excitation-inhibition balance in the M2-SC pathway may be a mechanism for dynamic modulation of collicular activity according to task demands.

**Fig. 6.**
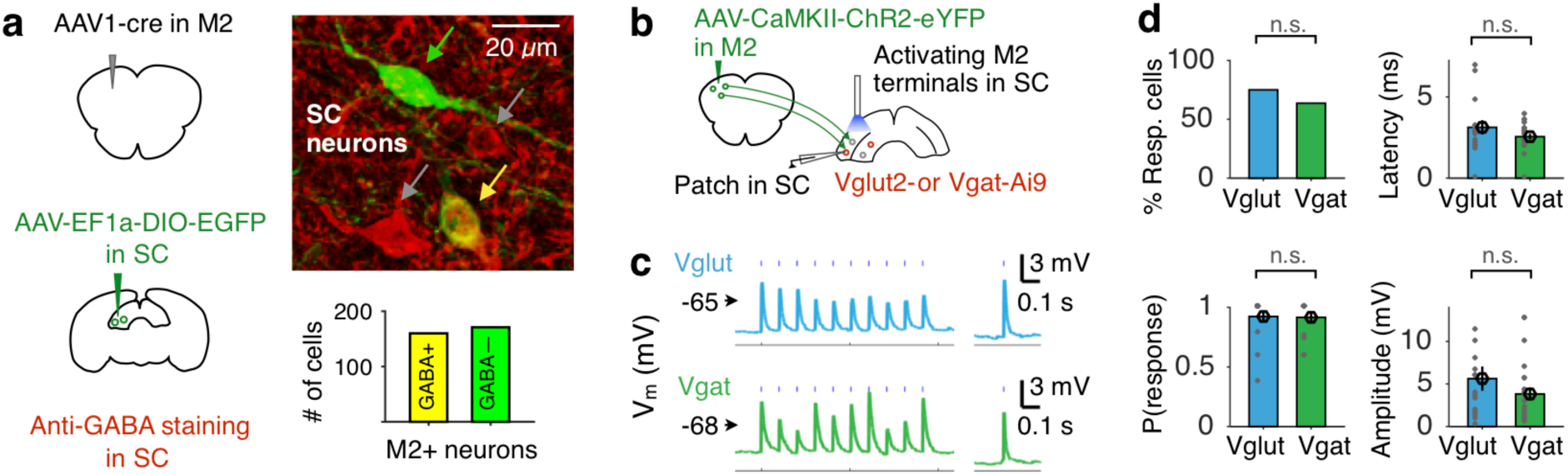
M2 terminals target SC excitatory and inhibitory neurons with comparable strength. **a**, Left, SC neurons with M2 inputs are labelled similar to 2**b** (green). GABA-ergic neurons in SC are labelled using anti-GABA staining (red). Right: green, yellow, and gray arrows point to SC neurons that are M2+ GABA-, M2+ GABA+, or M2-GABA+, respectively. 160 out of 331 M2+ SC neurons are GABA+; data from 3 mice. **b**, Design for cell-type-specific whole-cell recording in SC brain slice during optogenetic activation of M2 terminals. **c**, Membrane potential (Vm) changes in example excitatory (Vglut) and inhibitory (Vgat) SC cells, following a train of 10 pulses or single pulse of photostimulation, indicated by vertical ticks above the traces. **d**, Percent of responsive neurons, response latency, reliability (probability of response), and amplitude for excitatory (Vglut, n = 24) and inhibitory (Vgat, n = 44) SC neurons upon M2 terminal activation. Dots, individual cells; error bars, s.e.m. across cells. n.s., not significant; chi-squared or permutation tests.

## DISCUSSION

Our results indicate a critical role of the M2-SC pathway in decision maintenance, calling for a broader consideration of motor planning circuitry to include the midbrain SC as a central component in planning and executing spatial decisions. Recent studies showed that preparatory activity is redundantly and robustly represented in M2 on both hemispheres^35^; maintained via recurrent excitation in the thalamocortical loop^11^; and transformed into a motor command and sent to the brainstem^12^. The SC is interconnected with all these regions, and gated by the basal ganglia output for initiation of drinking behavior^31^. Although SC is well-situated to be a pivotal hub in the motor planning circuit, testing causal contributions of specific pathways without perturbing other input or output areas becomes a challenge. We combined projection-based imaging (Fig. 3,4) and inactivation (Fig. 5) methods with a psychometric delayed discrimination task (Fig. 1) to reveal the preferential involvement of the M2-SC pathway in decision-related processes during motor planning. M2 forms complex networks with multiple subcortical regions that have been shown to be involved in motor planning^7,11–13^. Such complexity can be even more puzzling, since the same M2 neurons project to both SC and brainstem^12^. Therefore, selectively inactivating the projection terminals in a specific target region, as shown in our results (Fig. 5), is key to determine the operation logic of this network, and should be applied to other downstream regions in future investigations.

In the primate oculomotor system, the frontal eye field (FEF) sends direct projections to SC^36,37^, and both FEF and SC project to the brainstem saccadic burst generator^38–40^, forming parallel premotor-brainstem and premotor-collicular-brainstem pathways underlying saccade planning and execution. Here, we combined different viral tracing tools to identify a putative SC lick region that receives M2 inputs and projects to the brainstem (Fig. 2a,b and Supplementary Fig. 2), with similar architectural features as the orienting circuit. Functionally, disrupting the information flow in the M2-SC pathway, without perturbing other M2 output projections, selectively impaired decision maintenance (Fig. 5), suggesting that contribution of the M2-SC-brainstem pathway cannot be fully compensated by direct M2-brainstem projections. These data are reminiscent of previous accounts in primates where the direct pathway from FEF to brainstem that bypasses the SC only plays a limited role in saccade generation^41^.

A growing body of work has observed convoluted relationships between task-relevant activity in a brain structure and the causal role of such activity during behavior^5,42,43^. The size of perturbation effects could positively^7,8^ or negatively^5^ correlate with the amount of choice information in an underlying area; and the specific timing of temporally-precise causal manipulation helps to set critical constraints on mechanistic explanations of decision-making^44,45^ and short-term memory^10,35^. Here, we found progressively stronger choice information in the M2-SC pathway that peaked during motor execution (Figs. 3,4), but the greatest perturbation effects occurred following delay-period inactivation, and not following response-period inactivation (Fig. 2). These data support a role of the M2-SC pathway in maintaining choice memory^46^. Consistent with this interpretation, comparing perturbation effects in task trials with short versus long delays revealed stronger impairment as delay duration increased (Fig. 5). Together, our results present empirical evidence for a distributed premotor-collicular network underlying decision-related processes in motor planning^5^, providing a critical foundation for future investigations of cortico-subcortical interaction during cognition and action.

## Acknowledgments

We thank J. Pan for technical support; E. Huang for help with the immunostaining experiments; J. C. Erlich for advice regarding the GLMM; A. T. Piet and C. D. Kopec for discussion regarding the dynamical attractor model literature; N. Li and K. Svoboda for discussion regarding the comparison between head-fixed mouse and freely-moving rat studies; X. H. Xu, X. K. Chen and Z. C. Guo for comments on the manuscript; and other members of the Xu lab for help and advice during all stages of the project.

## Funding

This work was supported by the “Strategic Priority Research Program” of the Chinese Academy of Sciences, Grant No. XDB32010000; National Natural Science Foundation of China, Grant No. 31571081; NSFC-ISF International Collaboration Research Project, Grant No. 31861143034; Key Research Program of Frontier Sciences, CAS, Grant No. QYZDB-SSW-SMC045; National Key R&D Program of China (grant No. 2017YFA0103900/2017YFA0103901); Shanghai Municipal Science and Technology Major Project, Grant No. 2018SHZDZX05; the Youth Thousand Talents Plan (to N.L.X.). C.A.D. is supported by the Simons Collaboration on the Global Brain Postdoctoral Fellowship and the CPSF-CAS Joint Foundation for Excellent Postdoctoral Fellows.

## Author contributions

C.A.D. and N.L.X. conceived the project and designed the experiments. C.A.D. developed the behavior and performed the optogenetic and imaging experiments. Y.P. performed the chemogenetic and immunostaining experiments. Y.P. and T.Z. performed the tracing experiments. Y.P., G.M., S.Z. performed the slice electrophysiology experiments. C.A.D. performed all the data analyses. C.A.D., N.L.X. and Y. P. wrote the manuscript.

## Competing interests

Authors declare no competing interests.

## Data and materials availability

All data is available in the main text or the supplementary materials. All the original behavioral, optogenetic, chemogenetic, electrophysiological, imaging and tracing data and analysis code are archived in the Institute of Neuroscience, Chinese Academy of Sciences, and can be obtained upon reasonable request via email to the correspondence authors.

## Methods

### Subjects

All animal use procedures were approved by the Animal Care and Use Committee of the Institute of Neuroscience, Chinese Academy of Sciences. Forty-one adult male mice were used for the experiments presented in this study. Of these, 11 C57BL/6J mice (SLAC) were used for chemogenetic inactivation experiments (8 in the experimental group and 3 in the control group). Nine Vgat-ires-Cre mice (Jackson Laboratory, JAX016962) were used for optogenetic inactivation experiments (6 for behavior testing and 3 for electrophysiological confirmation of optogenetic effect). Four mice (3 C57BL/6J and 1 Rbp4-Cre, MMRRC 031125) were used for two-photon calcium imaging experiments. Six mice (2 Vglut2-ires-Ai9, JAX016963 crossed with JAX007909; 4 Vgat-ires-Ai9, JAX016962 crossed with JAX007905) were used for whole-cell recordings in brain slices. Eleven mice were used for anatomical tracing and immunohistochemstry: 4 Rbp4-Cre mice were used for data in Fig. 2a; 2 C57BL/6J mice were used for data in Fig. 2b; 3 C57BL/6J mice were used for data in Fig. 6a; 2 C57BL/6J mice were used for data in Supplementary Fig. 2.

Mice were group-housed (<6 mice/cage) in a 12h reverse light:dark cycle, and all experimental procedures were conducted during the dark phase. Mice were water deprived before the start of behavioral training. On training days, mice received all their water inside training boxes. Supplementary water was provided for mice who could not maintain a stable body weight from task-related water reward.

### Behavior

In preparation for head-fixed behavioral training, a custom-made head-plate was implanted on each mouse during virus infection surgeries. The behavioral apparatus has been described previously^47^. In the final stage of the behavior (see training procedure below), mice were presented with auditory click trains of different rates (6 stimuli, log spaced between 20 and 125 Hz, 0.5 s). Click rates lower than 50 clicks/s indicated water reward from the left lick port; while higher click rates indicated reward from the right. Stimulus-response mapping was counterbalanced across animals. Each click was a 10 kHz pure tone lasting for 5 ms. Mice needed to discriminate and categorize the sensory cues, form and maintain a motor plan during a variable delay period (randomly sampled from a uniform distribution between 0.3 and 1.5 s), and eventually respond by licking one of the two lick ports after a “go” cue (pure tone, 5 kHz, 40 ms). Trials with premature licking before the “go” cue were terminated immediately, marked as violations, and excluded from further analyses. For non-violation trials, mice were allowed to make their responses within a 3-s window after the “go” cue. Failure to respond within the 3-s window was considered a “miss” trial. Response time (RT) was defined as time between the onset of the “go” cue and animals’ first lick after the “go” cue. Correct responses resulted in ∼6 μl of water reward. Error responses were followed by an “error” sound (pure tone, 20 kHz, 1 s), no reward, and a 2-6 s timeout. All trials ended with an inter-trial interval (ITI) of 2.5-3.5 s.

The general training procedure was inspired by methods described by Guo and colleagues^48^. In the first behavioral session, mice received water reward by licking either lick port. In the next 1-2 sessions, the rewarded port alternated between the left and right lick port after 3 rewards on each side, encouraging the mice to explore both lick ports. Each rewarded lick triggered an auditory stimulus (click trains, 20 or 125 Hz) associated with that side, followed by a 0.3 s delay, the “go” cue, and finally the water reward. The stimulus-response mapping remained constant throughout the training process.

After operant conditioning sessions, mice started two-alternative-forced-choice (2AFC) training and learned to actively discriminate two rates of auditory click trains (20 versus 125 Hz). On each trial, a 0.5-s stimulus was played, followed by a short variable delay (0.3-0.4 s), the “go” cue, and a 3-s response window during which mice’s licks were either rewarded or punished. The goal of this training stage was to establish the stimulus-response association; mice were not punished for licking before the “go” cue. We used three methods to facilitate learning. First, during the first 2AFC training session, mice were rewarded for correcting their mistakes within a 1-s window (termed as “grace period”) immediately after error responses. Second, error responses were followed by a 2-8 s timeout, during which continued licks to the wrong side reinitiated the timeout period. Third, following each error trial, the same type of trial was repeated once until mice made a correct response on that side.

Mice that reached criteria of >85% correct on the basic 2AFC task were then trained to withhold licking before the “go” cue. Any lick before the “go” cue (either during the stimulus or delay period) triggered a warning sound (noise stimuli, 1-5 kHz, 0.1 s) and a timeout, during which each additional lick resulted in the same warning sound and reinitiated the timeout period. These trials were terminated immediately and did not lead to any reward. For mice with high performance and low violation rates, delay duration was gradually increased to reach the final range (0.3-1.5 s). In the last phase of behavioral training, trials with intermediate click rates (log spaced between 20 and 125 Hz) were interleaved to test animals’ psychometric performance.

Mice trained on the final stage of the task were used for behavioral characterization and chemogenetic manipulation experiments. For optogenetic inactivation experiments, to ensure that all sub-trial inactivation conditions have the same duration for laser stimulation (0.5 s), mice were tested on a modified version of the behavior where the auditory stimulus period and the delay period both lasted 0.5 s instead of a variable delay period as in the original design. For two-photon calcium imaging experiments, we did not include trials with intermediate click rates due to limited trial numbers in imaging sessions.

### Surgery and virus injection

During surgery, mice were anesthetized with 1-2% isoflurane; and their body temperature was monitored and maintained at 37 °C using a heat-pad. Mice were placed in a stereotax with earbars. The skull was cleaned of blood and tissue, and leveled in the anterior-posterior (AP) direction so that the depth difference between bregma and lamda was within 50 µm. The locations of craniotomies for M2 and SC were marked on the skull using the stereotax. After craniotomies were performed, viral solution was injected into bilateral M2 (+2.5 mm AP, ±1.5 mm ML from bregma, 0.5 mm below brain surface) or bilateral SC (−3.5 mm AP, ±1.4 mm ML, +2.3 mm DV from bregma) over the course of ∼20 min/0.2 µL injection. The injection system comprised of a custom-made glass pipette (20-30 μm O.D. at the tip; Drummond Scientific, Wiretrol II Capillary Microdispenser) back-filled with mineral oil, and a fitted plunger inserted into the pipette to load or dispense viral solution. The plunger was controlled by a hydraulic manipulator (Narashige, MO-10); and the injection pipette was advanced into the brain using a Sutter MP-225 micromanipulator. After each injection, the glass pipette was left in the brain for 10 min before it was slowly withdrawn, to prevent backflow. Craniotomies were then covered with Kwik-Cast (World Precision Instruments), after which a thin layer of instant adhesive (Loctite 495) was applied onto the remaining skull. Finally, a custom-made head-plate was placed on the skull and cemented in place with an appropriate amount of dental acrylic. Post-surgery mice were allowed to recover for at least 3 days before water restriction.

### Chemogenetic inactivation

We used a DREADD (designer receptor exclusively activated by designer drug) based method^27^ to inactivate M2 (Supplementary Fig. 1), or M2 terminals in SC (Fig. 5). AAV2/9-Syn-hM4D(Gi)-mCherry (0.2 µL/site; titer: ∼10^13^ genomes/mL; Shanghai Taitool Bioscience Co. Ltd) was injected into bilateral M2. For chemogenetic inactivation of M2 terminals in SC, bilateral M2 neurons were infected with virus and bilateral SC were implanted with cannulae for local infusion. Guide cannula (22 AWG, PlasticsOne, VA) and dummy cannula (same length) were implanted into the brain at a 30-degree angle on each side until the final injector tip (32 AWG, extended 0.5-1 mm beyond the tip of guide cannula) reached −3.5 mm AP, ±1.4 mm ML, and +2.3 mm DV from bregma.

After at least two months of virus expression, we conducted intraperitoneal (IP) injection or cannula infusion of Clozapine-N-Oxide (CNO, Sigma) to inactivate infected neurons or terminals before behavioral testing. CNO was dissolved in saline (0.9% NaCl solution) to a stocking solution of 20 mg/mL; and stored as aliquots at −20 °C. For IP experiments, mice were briefly anesthetized with isoflurane, and 100 µL of saline or CNO (1-2 mg/kg) was injected 30 min before each behavioral session. For cannula infusion experiments, mice were either anesthetized with 1-2% isoflurane or head-fixed via the implanted head-plate during the infusion process. Dummy cannulae were removed and cleaned with 75% ethanol. Injectors were placed into guide cannulae and extended 0.5-1 mm past the end of the guide. A Hamilton syringe (0.5 µL) connected via tubing to the injector was used to infuse 0.2 µL of saline or CNO (5-10 µg/µL) into each brain area over the course of ∼30 s. The injector was left in the brain for 4 min to allow diffusion before removal. For bilateral infusions, the starting side was counterbalanced across sessions. After 4 min, cleaned and rinsed dummies were placed back into the guide cannulae, and mice were removed from isoflurane or head-fixation. After 30 min back in their home cages, mice were placed into the behavioral box for testing. Each saline control experiment with stable performance was followed by a CNO experiment in the same animal. The effect of inactivation was quantified as the difference in task performance between the CNO session and the saline control session the day before.

To control for the non-specific effects of CNO IP injection or infusion, we used a separate group of control mice (n = 3) where M2 neurons were infected with AAV2/9-Syn-mCherry (0.2 µL/site; titer: ∼10^13^ genomes/mL), and bilateral cannulae were implanted in the SC. All procedures were the same for the experimental and control groups other than the difference in virus infection. We did not observe any significant changes in performance in the control animals (Supplementary Fig. 1i-l and Supplementary Fig. 6e-g), suggesting that the chemogenetic inactivation results we observed in the experimental animals were not due to the non-specific effects of CNO injection, but due to the inactivation of neural activity.

### Analysis of behavior and chemogenetic inactivation data

To characterize animals’ baseline psychometric performance (Fig. 1d), we analyzed saline control sessions and non-photostimulation control trials. We combined data across sessions for each mouse and fit that data with a 4-parameter sigmoid function using Matlab’s nlinfit. The equation for the 4-parameter sigmoid function is as follows:

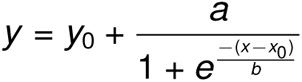

For these fits, y is ‘P(lick right)’, and x is ‘log(click rates)’ on each trial. The four parameters to be fit are: x_0_, the inflection point of the sigmoid; b, the slope of the sigmoid; y_0_, the lower bound of ‘P(lick right)’; and a+y_0_, the upper bound of ‘P(lick right)’. Psychometric curves for individual animals were plotted as thin gray lines (Fig. 1d). We then combined trials across all sessions from all animals, and generated a mean psychometric curve (black thick line, Fig. 1d). A similar fitting process was used to generate the psychometric curves for the saline versus CNO sessions in Fig. 5b,d and Supplementary Fig. 1c,k.

To quantify the effect of M2 soma inhibition (Supplementary Fig. 1) and M2 terminal inhibition (Fig. 5) across sessions, we calculated the performance difference between each pair of CNO and saline control sessions. Changes in error rate, violation rate, and miss rate were estimated using the means in each pair of sessions, and changes in RT were estimated using the medians in each pair of sessions. Nonparametric bootstrap procedures or permutation tests were then used to compute significance values across all sessions (shuffled 5000 times).

To investigate how M2 terminal inhibition affected error rates in different trial types, a repeated-measure two-way ANOVA was used to test the within-session main effect of difficulty (easy trials versus hard trials), delay duration (300-900 ms versus 900-1500 ms), and interaction of difficulty and delay on error increase due to inactivation. We also reported paired statistics computed using a permutation test (Fig. 5e).

### Optogenetic inactivation

For optogenetic inactivation of M2 or SC neurons, viral solution containing AAV2/9-Syn-FLEX-ChrimsonR-tdTomato (0.2 µL/site; titer: ∼10^13^ genomes/mL; Shanghai Taitool Bioscience Co. Ltd) was injected into bilateral M2 (+2.5 mm AP, ±1.5 mm ML from bregma, 0.5 mm below brain surface) and bilateral SC (−3.5 mm AP, ±1.4 mm ML, +2.3 mm DV from bregma) of 6 Vgat-IRES-Cre mice. After virus injection, four optic fibers with ceramic ferrule (200 µm in diameter, 0.37 NA, http://www.newdoon.com) were implanted in bilateral M2 (2 mm in length) and bilateral SC (3 mm in length) for a within-subject comparison of inactivated areas. The tips of optical fibers in M2 were pressed against dura; and the tips of optical fibers in SC were positioned 200 µm above the center of SC virus infection site (Supplementary Fig. 3c). Virus expression was allowed to develop for at least 2 months before behavioral testing began.

For each inactivation session, animals’ implanted fibers were connected to a red (635 nm) diode-pumped solid state laser (DPSS; Shanghai Laser & Optics Century Co., Ltd.) via an external optical fiber (200 µm in diameter, 0.37 NA). Laser power at the end of the external fiber was measured with a laser power meter (Sanwa, Mobiken series, LP1) and was adjusted using pulse-width modulation (PWM) to meet experimental requirement before each session. We used *in vivo* electrophysiology to search for stimulation parameters for complete inhibition with minimal rebound and minimal residual effect (see next section). For M2, the optimal parameters were 10 Hz laser pulses with 2 ms pulse width at 2 mW laser power measured at the tip of the external fiber (higher frequencies resulted in residual inhibition; Supplementary Fig. 3d). For SC, the optimal parameters were 20 Hz laser pulses with 2 ms pulse width at 2 mW laser power (lower frequencies resulted in incomplete inhibition; Supplementary Fig. 3e). For simultaneous inactivation of M2 and SC, we used a splitting optical fiber to target both regions at the same time. To ensure the temporal resolution of optogenetic inactivation, we used the lower stimulation parameters (10 Hz, 2 ms pulse width, 2 mW per fiber). Therefore, the simultaneous inactivation corresponded to complete inhibition of M2 and partial inhibition of SC, and can only be directly compared to M2 inhibition alone. Laser illumination occurred on 30% randomly chosen trials in each behavioral session. Different optogenetic conditions (stimulus, delay, response period, or inter-trial interval, 500 ms each) were randomly interleaved for all sessions to control for behavioral fluctuations across days.

The method of using GABAergic activation to silence local activity is well-established in cortex. However, subcortical structures such as the SC contain local GABAergic interneurons as well as GABAergic projection neurons, some of which project to the contralateral SC^16,49^. Therefore, the network effect of GABAergic activation in the SC may be more complex. Here, we used *in vivo* electrophysiology to confirm effective and temporarily-precise silencing of local SC activity (Fig 2d and Supplementary Fig. 3e). In addition, we observed an ipsilateral bias after unilateral SC inactivation, consistent with a more dominant effect of silencing local activity. Future experiments that selectively label GABAergic projection neurons versus local interneurons are needed to test their differential roles during motor planning.

### Electrophysiological verification of optogenetic effects

To measure the effects of optogenetic inactivation on neural activity, acute recordings of infected M2 or SC neurons were performed in awake mice using custom made optrodes (Supplementary Fig. 3d,e). A sharp tungsten electrode (0.5 or 1.0 MΩ, World Precision Instruments, Inc.) was glued to a stripped optical fiber (200 µm in diameter, 0.37 NA), with the tip of the optical fiber 200-300 µm above the tip of the tungsten electrode to parallel conditions in optogenetic inactivation experiments. The optrode was then threaded into a glass tubing for protection; glued in position with epoxy so that <1 cm of the optrode protruded from the glass tubing. During acute recording sessions, the optrode was advanced to the center of the infected area in awake head-fixed mice. For each neuron tested, baseline neural activity was recorded for 4 s, followed by 0.5 or 3 s of laser stimulation, and another period of post-stimulation recording, repeated for 20 times. We systematically tested different laser power, laser pulse width and pulse frequency to find optimal stimulation parameters for temporally-precise inactivation of M2 and SC neurons.

### Analysis of optogenetic inactivation data

To estimate the effect size of the ipsilateral bias due to optogenetic inactivation, we compared P(lick ipsi) on inactivation trials and control trials from the same sessions. For each session, we calculated mean P(lick ipsi) on control trials, and subtracted that mean value from the P(lick ipsi) of individual inactivation trials. After obtaining the normalized ipsilateral bias change for trials in each session, we concatenated trials across all sessions and all mice, and computed the mean and s.e.m. of delta ipsilateral bias across trials.

To visualize the effects of optogenetic inactivation on animals’ psychometric performance, we fit a 4-parameter sigmoid function to control and inactivation trials across all sessions for each inactivation epoch (Fig. 2, g,j,m), similar to the fitting process used to generate Fig. 1d. These fits are for visualization only. All statistics reported for optogenetic inactivation experiments were computed using generalized linear mixed models (GLMM) as implemented in the function ‘glmer’ in package ‘lme4’ (R). For unilateral M2 (Fig. 1g) or SC (Fig. 1j) inactivations, we fit a mixed-effects model where mice’s choice on each trial was a logistic function of log(click rates) and optogenetic inactivation epoch (ITI, stimulus, delay, response period, or no inactivation) as fixed effects. The mouse and an interaction of mouse, log(click rates), and optogenetic inactivation time were modeled as within-subject random effects. The statistics reported for M2 or SC inactivations were the fixed effect *P* values for the ipsilateral bias change due to optogenetic inactivation in that specific epoch compared to the no inactivation condition. We then conducted multiple comparisons (‘mcp’ in ‘multcomp’, R) of the GLMM results to simultaneously test whether the delay period inactivation effect was significantly different from the stimulus period effect and the response period effect. Adjusted *P* values from this single-step method were reported.

To compare inactivation results in different brain regions at different behavioral epochs, we used a GLMM to fit combined inactivation trials from M2 inactivation, SC inactivation, and simultaneous M2 and SC inactivation during the ITI, stimulus, delay, and response period (Model 1, Supplementary Fig. 4). In this model, mice’s choice on each trial was a logistic function of log(click rates), optogenetic inactivation epoch, and optogenetic inactivation region as fixed effects and within-subject random effects. We reported adjusted *P* values of the multiple comparison results that tested if the stimulus, delay, response period effects were significantly different from each other; and if the M2, SC, or M2+SC inactivation effects were significantly different from each other.

To test how optogenetic inactivation affected hard versus easy trials differently, we included a three-way interaction term (epoch : region : difficulty) to Model 1 described above (Model 2, Supplementary Fig. 4). Difficulty was defined as a quadratic function of the normalized log(click rates). Defining difficulty as 1 minus the absolute value of the normalized log(click rates) did not change the main effects. We demonstrated that Model 2 was a significantly better fit to the data than Model 1 using a likelihood ratio test (‘anova’, R). Fixed effect *P* values for the interaction terms were reported.

### Two-photon calcium imaging

Two methods were used to selectively label M2 neurons that project to the SC. For three C57BL/6J mice, AAV2-Retro-Cre (titer: ∼10^13^ genomes/mL; diluted threefold in saline; 0.2 µL; Shanghai Taitool Bioscience Co. Ltd) was injected into the left SC (−3.5 mm AP, −1.4 mm ML, +2.3 mm DV from bregma). After behavioral training, a circular craniotomy (∼2.5 mm in diameter) was made above the left M2 (centered at +2.5 mm AP, −1.5 mm ML from bregma), and AAV2/9-Syn-FLEX-GCaMP6s (titer: ∼10^13^ genomes/mL; diluted threefold in saline; Shanghai Taitool Bioscience Co. Ltd) was injected at 3-5 locations (50 nl per site) at a depth of 0.5-0.6 mm from brain surface. With this method, only SC-projecting M2 neurons would express GCaMP virus; and both apical dendritic trunks and somas were imaged at different sessions. For one Rbp4-Cre mouse, red RetroBeads (diluted fivefold in saline; 0.2 µL; Lumafluor) were injected into the left SC, and AAV2/9-Syn-FLEX-GCaMP6s (titer: ∼10^13^ genomes/mL; diluted threefold in saline; Shanghai Taitool Bioscience Co. Ltd) was injected into the left M2. With this method, only GCaMP-expressing M2 somas with beads co-localization were identified as SC-projecting M2 neurons. Custom-designed imaging window (Fig. 3a) was constructed from an outer steel ring, a cannula tube (0.6 mm height), and one layer of microscope coverglass (2.3 mm in diameter) glued to the outer surface of the cannula tube. During the first surgery, virus or RetroBead was injected into the SC, and the location of M2 craniotomy was marked on the skull before a custom-made head-plate was implanted. After behavioral training, mice underwent the second surgery, where M2 craniotomy and GCaMP viral injection were performed before fixing the imaging window to the skull using dental acrylic. Water restriction started 3-5 days after each surgery.

Imaging experiments started 3-4 weeks after GCaMP viral infection. Images were acquired using a custom-built two-photon microscope (http://openwiki.janelia.org/wiki/display/shareddesigns/MIMMS), with a resonant galvanometer (Thorlabs; 16 kHz line rate, bidirectional). GCaMP6s was excited using a Ti-Sapphire laser (Coherent) at 920 nm, and the average laser power for imaging SC-projecting M2 somas and apical trunks was ∼160 mW and ∼120 mW respectively. Emitted fluorescence was isolated using a bandpass filter (525/50, Semrock). The objective was a 16× water immersion lens (Nikon, 0.8 NA, 3 mm working distance). The field of view was 300 by 300 um (512 x 512 pixels), imaged at ∼30 frames/s. The entire microscope was enclosed in a custom-designed sound attenuation box, and the system was controlled using ScanImage (http://scanimage.org). Images were acquired continuously for the entire session. To identify the M2 neurons retrogradely labeled with Retrobead, a small image stack was acquired around the imaging depth at the end of each functional imaging session (laser wavelength = 840 nm).

### Analysis of calcium imaging data

Brain motion was corrected using a cross-correlation-based registration algorithm^50^. Regions of interest (ROIs) were manually selected based on identifiable cell bodies and dendritic apical trunks. The boundaries of the ROIs were determined using mean and maximum fluorescence intensity across frames of all trials. The fluorescence (F) time series of each ROI was estimated by averaging all the pixels within the ROI. Slow timescale fluorescence changes were corrected by determining the distribution of fluorescence values over 400 frames (∼14 s) around each time point and subtracting the eighth percentile value^51^. ΔF/F_0_ was then calculated for each ROI as (F - F_0_) / F_0_, where F_0_ is the mode of F over the entire session after slow timescale fluorescence correction. For each trial, averaged fluorescence signal before sound onset was subtracted from activity at each frame.

Due to the variable delay and response durations across trials, we could not rely on any one time point alignment to fairly compare neural selectivity across all behavioral epochs. Instead, we first extracted and averaged neural responses during the relevant time window for each behavioral epoch per trial (Fig. 3e). For variable delay durations, activity within 500 ms before the “go” cue was taken for each trial. Response period activity was extracted from the entire response time, which varied across trials. Using these epoch-averaged fluorescence values (Fig. 3f), single-neuron and population-level analyses were conducted for all behavioral epochs across trials.

To quantify single neuron selectivity, we conducted receiver operating characteristic (ROC)^34^ analysis on responses in each behavioral epoch, and calculated the area under the ROC curve (AUC) to indicate discrimination between trial types. Correct and error trials were included to calculate choice selectivity (lick left versus lick right trials) and sensory selectivity (high rate versus low rate trials) for each neuron. To compute the significance of AUC values, we shuffled the trial labels 1000 times to obtain a distributions of AUC values for shuffled data. For a given neuron, if the sensory or choice AUC was outside the 99% confidence interval of the shuffled data during any behavioral epoch, that cell was labeled as significantly side-selective (Fig. 3b,c).

To characterize the changes in choice AUC values across behavioral epochs (Fig. 3g and Supplementary Fig. 5b,c), we converted each neuron’s choice AUC value to range between 0 and 1, where 0 represents strong preference for ipsilateral choices, 1 represents strong preference for contralateral choices, and 0.5 represents no choice preference. For each behavioral epoch, the histogram and gaussian fit to the distribution of converted choice AUC values for all individual neurons were plotted for one example animal (Fig. 3g) or for all animals (Supplementary Fig. 5b,c). Individually significant AUC values (*P* < 0.05) were marked in black. For each pair of epochs (sound versus delay, delay versus response, response versus lick), a non-parametric permutation test (shuffled 5000 times) was used to compute statistical difference between distributions of choice AUC values (*P* < 0.01 for all pairs in the example animal, *P* < 10^-3^ for all pairs for data across all animals).

To measure the amount of choice or sensory-related information in simultaneously imaged neural populations (Fig. 4a-c), we performed a series of cross-validated linear classification analyses (support vector machine, SVM, ‘fitcsvm’, Matlab). For each session, simultaneously imaged calcium signals were arranged in a M×N×T matrix, where M is the number of trials, N is the number of neurons, and T is the number of time points (task epochs). For a given trial and one task epoch, the vector of N neurons’ ΔF/F_0_ values represents the population response vector for that condition (Fig. 4a). To decode choice or sensory-related information, we randomly sampled 90% of imaged trials as the training set, and the remaining 10% of trials as the test set. The training set was used to compute the linear hyperplane that optimally separated the population response vectors corresponding to lick-left versus lick-right trials (choice decoding) or high-rate versus low-rate trials (sensory decoding). The number of training trials were balanced between conditions to avoid bias in favor of one condition over the other. Performance was calculated as the fraction of correct classifications of the test trials. Decoders were trained and tested independently for each behavioral epoch. We repeated the resampling process 100 times, and computed the mean and standard error of decoding accuracy across the 100 resampling iterations. To test if one session’s decoding accuracy was significantly higher than chance (0.5), or if choice decoding was significantly better than sensory decoding, we used bootstrap or permutation tests.

To investigate the relationship between animals’ response time (RT) and the amount of choice information during the delay on a trial-by-trial basis, we used leave-one-out cross-validation (Fig. 4d,e). For each session with M trials, the training and testing procedure was repeated M times. For each iteration, we trained the decoder on all but one trial (balanced between conditions); and tested the classification performance (0 or 1) for that one test trial. We then binned trials based on RT (0 - 2.5 s), and calculated the mean choice decoding accuracy for each 20 percentile of RT per session (Fig. 4e, right). The relationship between decoding accuracy and RT was quantified using linear correlation. Because delay duration and delay-period neural activity were both negatively correlated with RT, we fit a linear mixed model (LMM, ‘fitlme’, Matlab) to simultaneously consider their contributions for predicting RT on a trial-by-trial basis. Animals’ RT on each trial was modeled as a linear combination of delay duration and the classification performance for that trial, both as fixed effects and as within-subject random effects. The LMM statistics reported were the fixed effect *P* values for predicting RT (delay duration or choice decoding accuracy). Including behavioral performance on each trial (correct or error) as a third fixed effect in the LMM did not change the main conclusions.

### Slice preparation and recording

To study how M2 activity modulates different types of SC neurons, we performed *in vitro* slice electrophysiology recording. We injected AAV-DJ-CaMKIIa-ChR2-EYFP virus into unilateral M2 (+2.5 mm AP, ±1.5 mm ML from bregma, 0.5 mm below brain surface, 0.5 µL/site; titer 1.81E14 v.g./ml; BrainVTA Co. Ltd) and recorded SC neurons’ responses in acute brain slices upon optogenetic stimulation of M2 terminals. Transgenic mice were used to label excitatory (Vglut2-ires-Ai9, JAX016963 crossed with JAX007909) and inhibitory (Vgat-ires-Ai9, JAX016962 crossed with JAX007905) neurons. Three to four weeks after virus injection, mice (∼10 weeks of age) were sacrificed and coronal brain slices (300 µm thickness) were acquired with vibratome (LEICA VT 1200s) in ice-cold recovery solution (containing 93 mM NMDG, 2.5 mM KCl, 1.2 mM NaH_2_PO_4_, 30 mM NaHCO_3_, 20 mM HEPES, 25 mM glucose, 5 mM Sodium ascorbate, 2 mM Thiourea, 3 mM Sodium pytuvate, 10 mM MgSO_4_, 0.5 mM CaCl_2_). Sections were immediately transferred to recovery solution at 32℃ for 5 min and then incubated in ACSF (containing 125 mM NaCl, 2.5 mM KCl, 1.25 mM NaH_2_PO_4_, 1 mM MgCl_2_, 2 mM CaCl_2_, 12.5 mM Glucose, 25 mM NaHCO_3_) at room temperature till recording.

To find regions with high density of M2 terminals and target tdTomato+ cells, we examined expression pattern in each brain slice under a fluorescence microscope (Olympus U-TV1 X). Borosilicate glass pipettes containing intracellular solution (135 mM K-gluconate, 5 mM KCl, 10 mM HEPES, 0.3 mM EGTA, 4 mM MgATP, 0.3 mM Na_2_GTP, 10 mM Na_2_-phosphocreatine) with impedance larger than 3MΩ were used to patch neurons. Signals were recorded with a MultiClamp 700B amplifier (Axon Instrument), under current clamp mode, filtered at 2 kHz and sampled at 10 kHz. Single light pulse (5 ms, 25mW power from the objective) or a train of ten pulses (10 Hz, each pulse 5 ms) were delivered to activate M2 terminals using a mercury arc lamp via the objective of the microscope.

### Analysis of slice recording data

To determine if an identified excitatory or inhibitory SC neuron is responsive to M2 terminal activation, we repeatedly delivered trains of 10-pulse photostimulation and calculated the rate of significant responses based on average EPSPs over repeated trials. Light evoked voltage changes (within 50 ms after light delivery) exceeding 3 s.d. of baseline voltage (over 1 s before stimulation) were considered significant transients; and cells with >40% significant transients were considered responsive. A chi-squared test was used to calculate statistical difference between the percentage of responsive inhibitory neurons versus excitatory neurons.

For responsive neurons, we repeatedly delivered single-pulse photostimulation to further characterize their response reliability, latency, and amplitude. Reliability was calculated as the probability of significantly responsive trials for each neuron. Response latency and amplitude were calculated using mean response traces averaged over all individual trials. Latency of EPSP corresponded to the amount of time between light stimulation onset and the first time point when voltage change exceeded 3 s.d. of baseline. Response amplitude corresponded to the maximum voltage change above baseline within 50 ms after light delivery. Permutation tests (shuffled 5000 times) were used to report statistical difference between two groups of SC neurons.

### Tracing experiments

All viruses used in tracing and immunostaining experiments were provided by Shanghai Taitool Bioscience Co. Ltd and had a titer of ∼10^13^ genomes/mL. To identify the part of SC downstream of M2 (Fig. 2a), we unilaterally injected AAV-Syn-FLEX-EGFP (0.2 µL/site) into M2 (+2.5 mm AP, ±1.5 mm ML from bregma, 0.5 mm below brain surface) of Rbp4-Cre mice, which constrained virus expression to M2 layer 5 neurons.

To label SC somas that receive M2 projections, we applied anterograde transsynaptic tracing (Fig. 2b) using one serotype of adeno-associated virus (AAV1)^32^. AAV1-hSyn-Cre (0.2 µL/site) was unilaterally injected into M2 of C57 mice, and AAV-hSyn-FLEX-tdTomato (0.2 µL/site) was injected into the ipsilateral SC (−3.5 mm AP, ±1.4 mm ML from bregma, 2.3 mm below bregma).

To compare the projections of M2 and SC in the brainstem, we simultaneously labeled M2 neurons and SC neurons downstream of M2 (Supplementary Fig. 2) with different fluorescence. We injected a mixture of virus containing AAV1-hSyn-Cre and AAV-hSyn-FLEX-tdTomato (volume ratio, 1:1, 0.1 µL/site in total) unilaterally into the M2 of C57 mice. In the ipsilateral SC, we also injected AAV-CAG-FLEX-EGFP (0.2 µL/site). Red and green terminals in the brainstem represent direct projections from M2 and projections from SC neurons downstream of M2, respectively.

For all experiments mentioned above, mice were sacrificed 4 weeks after virus injection. Brain slices were imaged using Olympus VS120 Virtual Slide Fluorescence Microscope.

### Immunostaining

To characterize the types of SC neurons that receive M2 projections, we combined anterograde transsynaptic viral tracing with immunohistochemistry (Fig. 6a). We injected AAV1-hSyn-Cre (50nL/site) unilaterally into M2 and AAV-EF1a-DIO-EGFP (0.2 µL/site) into the ipsilateral SC. SC neurons downstream of M2 were thus labeled with green fluorescence. Four weeks after virus injection, mice were perfused using 4% paraformaldehyde (PFA). Brains were incubated in PBS solution containing 30% sucrose overnight at room temperature and coronally sectioned with a cryostat (20 µm). Slices containing SC neurons were collected and underwent immunostaining.

After incubated with blocking buffer (PBS containing 5% Bovine Serum Albumin and 0.3% Triton X-00) for 2 hours at 37 °C, sections were incubated with the primary anti-GABA antibody (rabbit, dilution 1:1000, A2052; Sigma) overnight at 4 °C. The primary antibody was then washed three times with PBS (15 min each) before adding the secondary antibody (donkey anti-rabbit, Alexa Fluor 594, dilution 1:1000, A-21207; ThermoFisher Scientific). Brain sections were incubated in the secondary antibody for 2 hours at 37 °C. Finally, the secondary antibody was washed 3 times (15 min each) with PBS and sections were mounted onto microscope slides with reagent containing DAPI to stain nuclei. Sections were imagined with Nikon A1 inverted confocal microscope (X20 objectives) and confocal images were analyzed with ImageJ.

To quantify the percentage of GABA+ cells (red) in total SC neurons that receive M2 projections (green), we sampled three slices (−3.28 to −4.04 mm AP from bregma) for each animal and manually counted EGFP labeled SC neurons. Among these SC neurons downstream of M2, we checked whether they were GABA immunoreactivity positive (red) to calculate the ratio of GABAergic versus non-GABAergic neurons.

### Histology

Histology was performed on all mice used for imaging, electrophysiology, chemogenetic and optogenetic experiments after data collection was completed. Animals were anesthetized and perfused with 4% PFA or with 10% formalin, and brains were removed and post-fixed overnight. Brains were then incubated with PBS solution containing 30% sucrose overnight. Frozen sections (50 µm) containing viral injection sites, cannula implantation tracks or optic fiber tracks were gathered and stained with DAPI to visualize nuclei. Pictures were taken with Olympus MacroView MVX10 fluorescent microscope.

For the chemogenetic inhibition experiments, six out of eight mice that underwent surgeries had cannulae placed correctly within the SC. These six mice then included for analyses in the M2 terminal inhibition experiment (Supplementary Fig. 6b). The two mice that had cannulae placed outside of the SC were only included for analyzing M2 soma inhibition experiment but excluded for the terminal inhibition experiment. For the optogenetic inhibition experiments, all six mice were included as the final dataset based on effective virus expression and correct optic fiber placements (Supplementary Fig. 3c).

---

**Supplementary Fig. 1.**
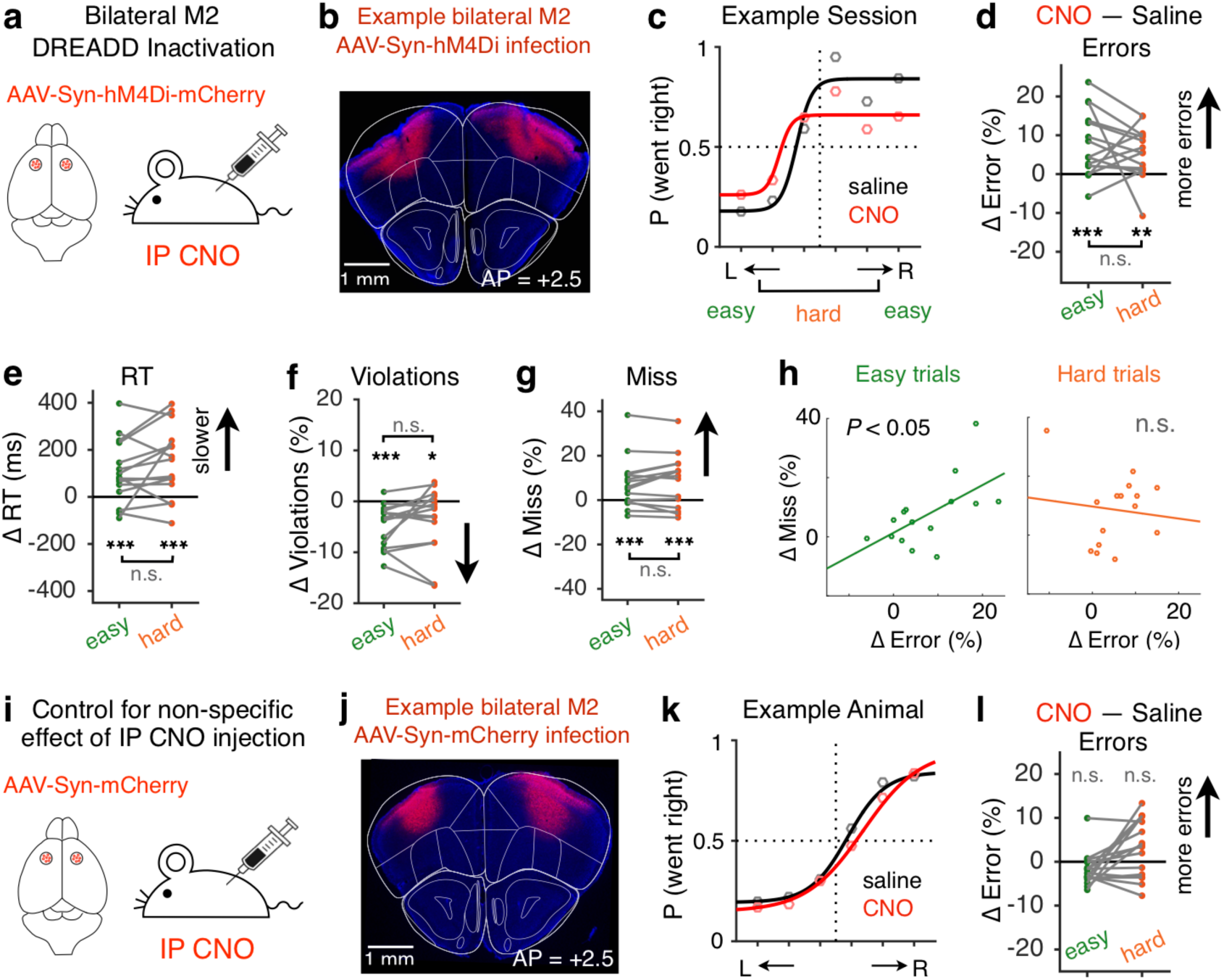
Chemogenetic inactivation of M2 during behavior. **a**, Design for bilateral M2 DREADD inactivation. **b**, Example histology of bilateral M2 infection for chemogenetic inactivation. **c**, Example M2 inactivation session (red) compared to saline control session (black) for one animal. Dots, data; lines, sigmoid fit. **d-g**, Difference in error rate (**d**), RT (**e**), violation rate (**f**), and miss rate (**g**) between M2 inactivation sessions (n = 15) and saline control sessions (n = 15) for easy (green) and hard (orange) trials. Gray lines connect data from the same session. \**P* < 0.05; \*\**P* < 0.01; \*\*\**P* < 10^-3^; n.s., not significant; bootstrap. **h**, Session-by-session correlation between changes in miss rate and error rate for easy (left) and hard (right) trials. **i-l**, Similar to **a-d**, for control animals (n = 16 sessions, 3 mice) with AAV-Syn-mCherry viral infection. IP injection of CNO itself did not result in increased error rates in control animals.

**Supplementary Fig. 2.**
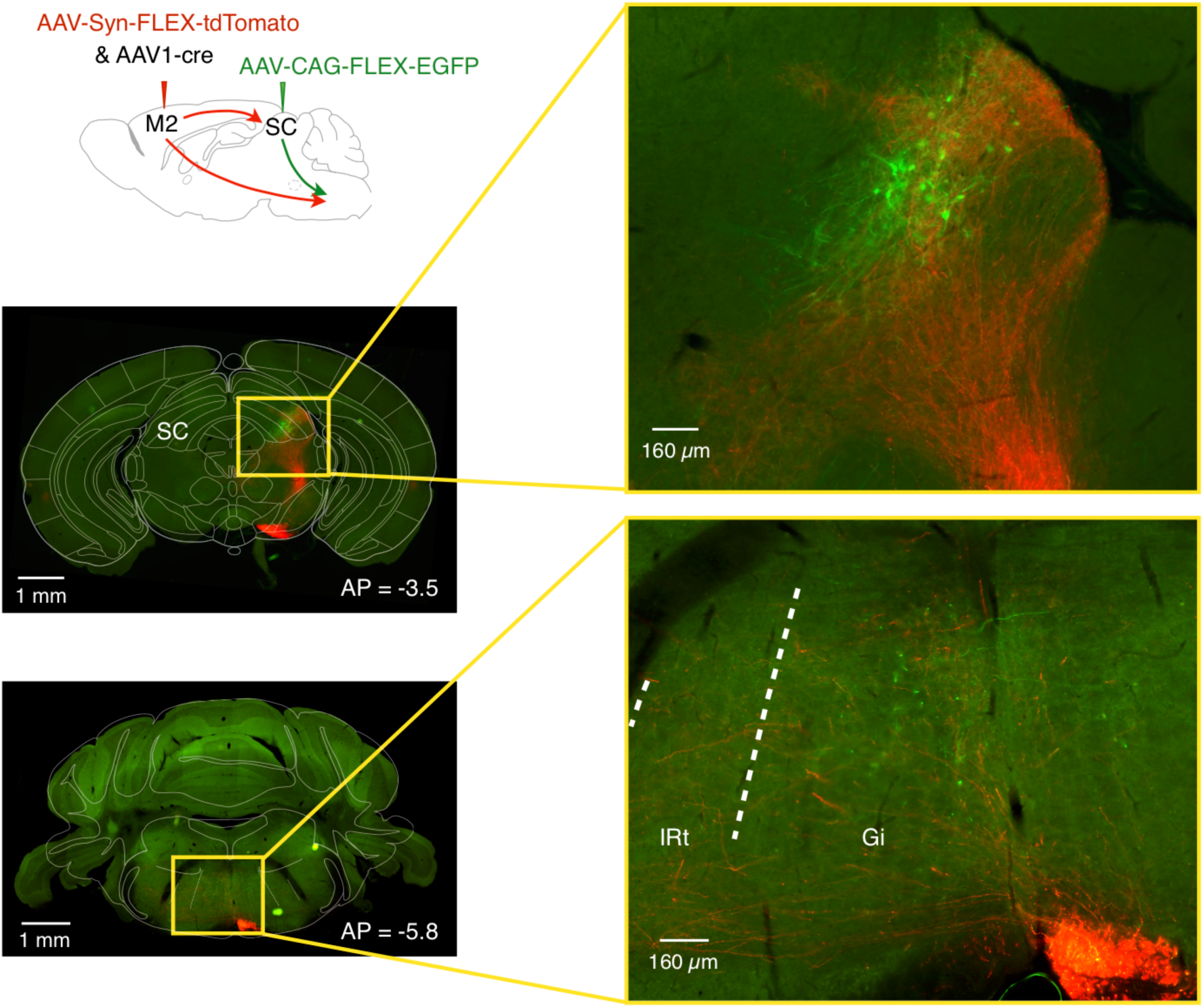
Descending projections from M2 and lateral SC. To examine the descending projections from M2 and SC to the brainstem, we used anterograde transsynaptic virus to simultaneously label M2 neurons (red) and SC neurons (green) downstream of M2. Red and green terminals in the brainstem represent direct projections from M2 and projections from SC neurons downstream of M2, respectively. These data suggest parallel descending pathways for generating licking responses. Gi, gigantocellular reticular nucleus. IRt, intermediate reticular formation.

**Supplementary Fig. 3.**
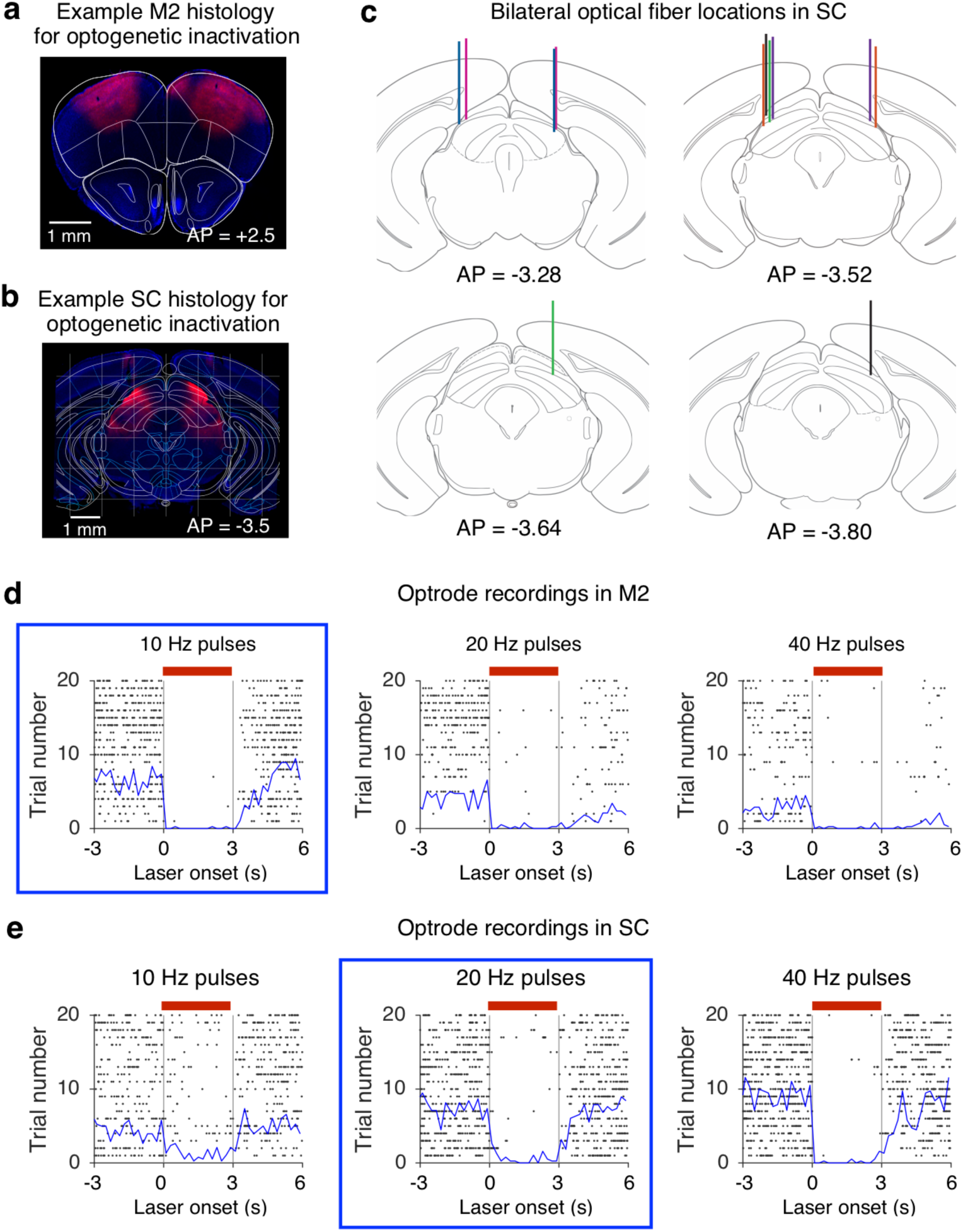
Optogenetic inactivation in M2 and SC. **a-b**, Example histology of bilateral M2 (**a**) and SC (**b**) infection for optogenetic inactivation. **c**, Optical fiber tracks for all mice (n = 6) used for SC optogenetic inactivation experiments. Bilateral fiber tracks from the same animal are labeled with the same color. We positioned the optical fibers to be 200-300 µm above the center of lateral SC virus infection site, −3.5 mm from bregma on the anterior-posterior (AP) axis. The actual fiber tracks were found in coronal sections ranging from −3.80 to −3.28 mm from bregma on the AP axis. **d**, Spike activities of an example M2 neuron under 3 types of optogenetic inactivation conditions. Spikes are aligned to laser onset over 20 repeated trials per condition. The red bar marks the laser illumination period (3 s), and the blue line plots the average firing rate as a function of time. In our study, the optimal laser pulse frequency for M2 inactivation is 10 Hz (left panel); higher pulse frequencies resulted in residual inhibition (middle and right panels). e, Spike activities of an example SC neuron under 3 types of optogenetic inactivation conditions, similar to (**d**). The optimal laser pulse frequency for SC inactivation is 20 Hz (middle panel); lower pulse frequencies resulted in incomplete inhibition (left panel).

**Supplementary Fig. 4.**
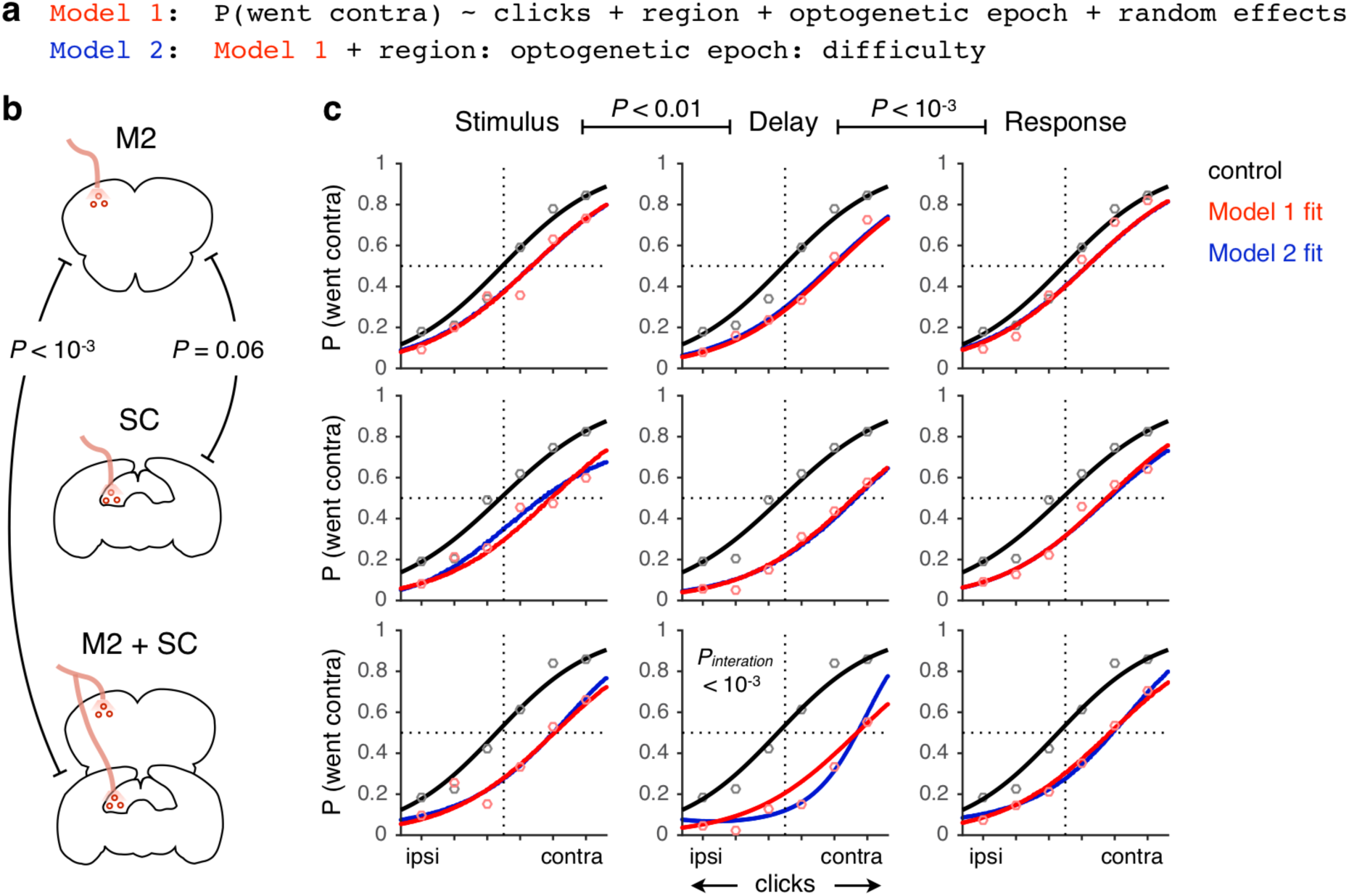
Simultaneous optogenetic inactivation of M2 and SC. **a**, Pseudo-formula for two GLMMs (logistic functions) used to fit inactivation data. Model 2 includes an extra three-way interaction term compared to Model 1. b, Schematic for optogenetic inactivation of unilateral M2, SC, or M2 and SC on the same hemisphere. c, Data (circles, 16391 trials from 36 sessions) and model fit (lines) for optogenetic experiments across different regions and epochs of inactivation. GLMM (model 1, red curves) followed by single-step multiple comparisons (corrected *P* values) reveals a preferential involvement of the cortico-collicular circuit during the delay; and simultaneous inactivation of M2 and SC resulted in a larger impairment than cortical inactivation alone. Model 2 (blue curves) is fit to test which experiment affects difficult trials more compared to easy trials. This interaction term is significant only for simultaneous M2 and SC inactivation during the delay period, and not significant for any other condition (*P*’s > 0.05).

**Supplementary Fig. 5.**
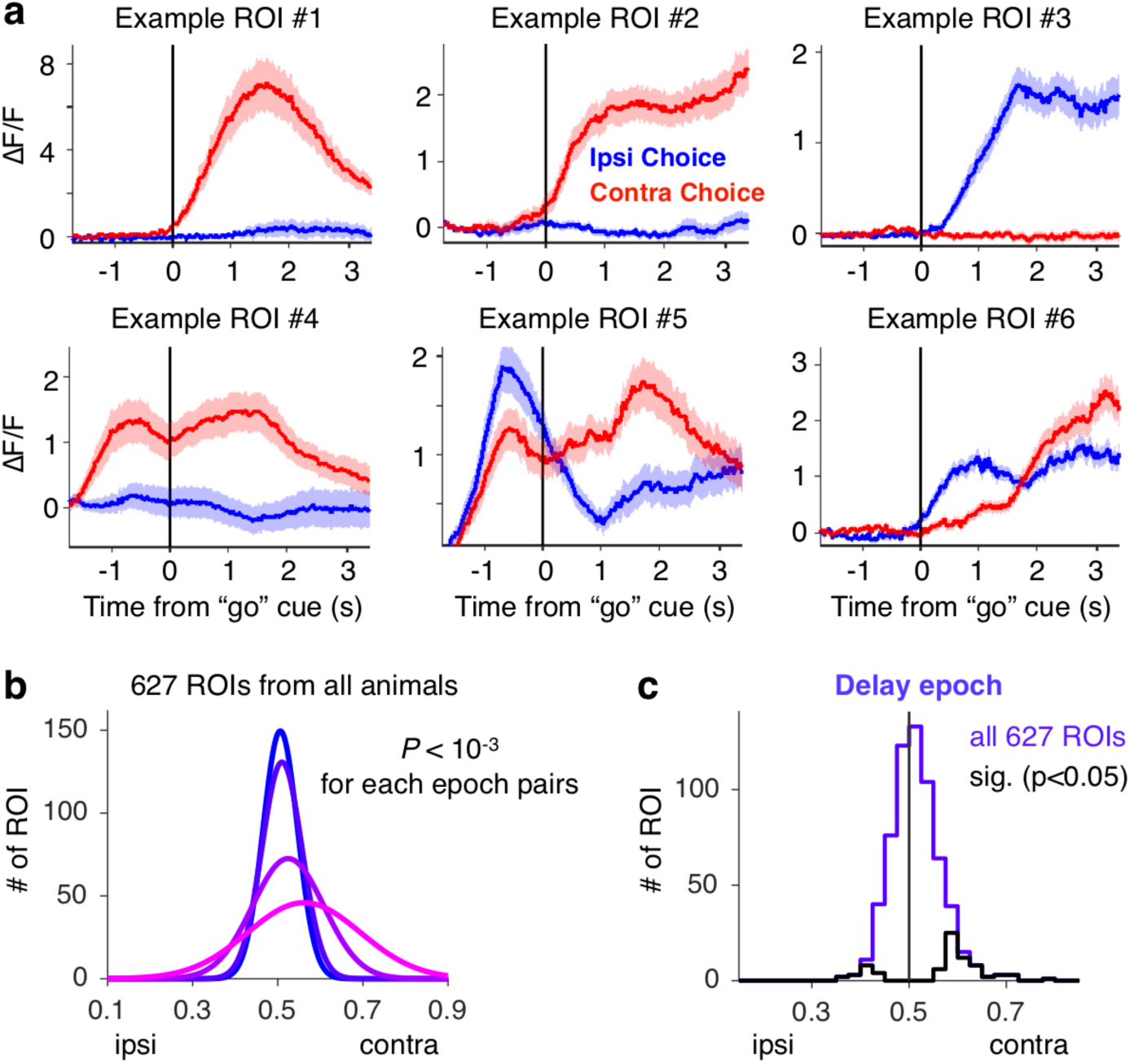
Single neuron properties of SC-projecting M2 neurons. **a**, Six example SC-projecting M2 neurons. Mean ± s.e.m. ΔF/F traces for contra (red) and ipsi (blue) choice trials, aligned to the “go” cue. **b**, Gaussian fits to distributions of choice AUC values of all the ROIs (4 animals) during the sound, delay, response, and lick epochs, similar to Fig. 3g. **c**, Histograms of choice AUC values of all the ROIs (4 animals) during the delay epoch. Black lines show individually significant neurons.

**Supplementary Fig. 6.**
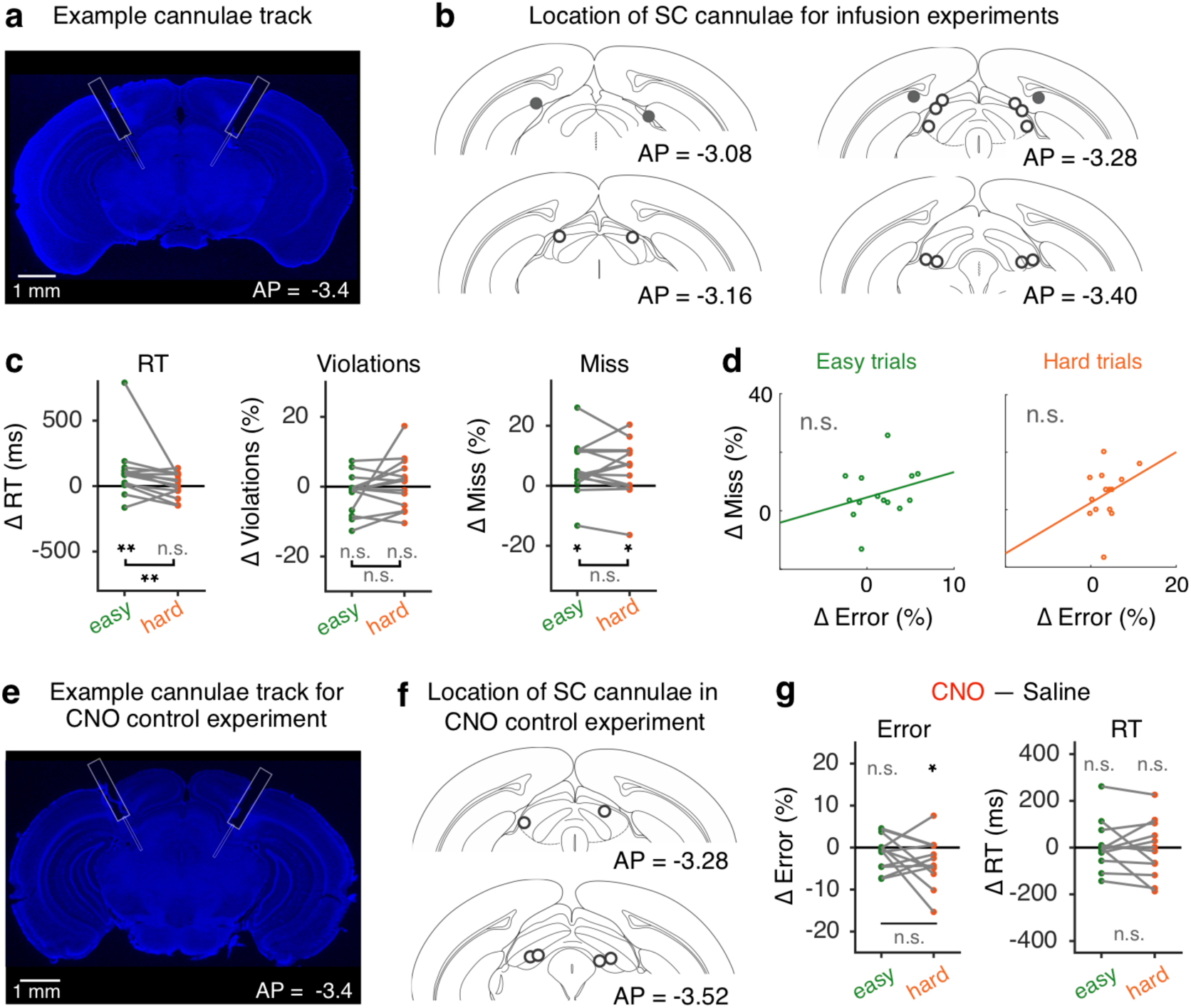
Pathway-specific inactivation of M2 terminals in SC. **a**, Example cannulae track for SC infusion experiment. **b**, Histology of bilateral SC cannulae placement for all mice implanted for M2 terminal inhibition experiments (n = 8 mice). Open and closed circles mark the locations of the injector tips that were within the SC or outside the SC, respectively. Six out of 8 mice that underwent surgeries had cannulae placed correctly within the SC, and were included for analyses in the M2 terminal inhibition experiment. **c**, Difference in RT (left), violation rate (middle), and miss rate (right) between M2 terminal inhibition sessions (n = 14) and saline control sessions (n = 14) for easy (green) and hard (orange) trials. Gray lines connect data from the same session. \**P* < 0.05; \*\**P* < 0.01; n.s., not significant; bootstrap. **d**, Session-by-session correlation between changes in miss rate and error rate for easy (left) and hard (right) trials after chemogenetic inactivation of M2 terminals in SC. **e-g**, Similar to **a-c**, for control animals (n = 12 sessions, 3 mice) with AAV-Syn-mCherry viral infection in M2 neurons.

## References

1. Funahashi, S., Bruce, C. J. & Goldman-Rakic, P. S. Neuronal activity related to saccadic eye movements in the monkey’s dorsolateral prefrontal cortex. J. Neurophysiol. 65, 1464–1483 (1991).

2. Erlich, J. C., Bialek, M. & Brody, C. D. A cortical substrate for memory-guided orienting in the rat. Neuron 72, 330–343 (2011).

3. Bruce, C. J. & Goldberg, M. E. Primate frontal eye fields. I. Single neurons discharging before saccades. J. Neurophysiol. 53, 603–635 (1985).

4. Dias, E. C. & Segraves, M. A. Muscimol-induced inactivation of monkey frontal eye field: effects on visually and memory-guided saccades. J. Neurophysiol. 81, 2191–2214 (1999).

5. Kopec, C. D., Erlich, J. C., Brunton, B. W., Deisseroth, K. & Brody, C. D. Cortical and Subcortical Contributions to Short-Term Memory for Orienting Movements. Neuron 88, 367–377 (2015).

6. Murakami, M., Shteingart, H., Loewenstein, Y. & Mainen, Z. F. Distinct Sources of Deterministic and Stochastic Components of Action Timing Decisions in Rodent Frontal Cortex. Neuron 94, 908–919.e7 (2017).

7. Li, N., Chen, T.-W., Guo, Z. V., Gerfen, C. R. & Svoboda, K. A motor cortex circuit for motor planning and movement. Nature 519, 51–56 (2015).

8. Guo, Z. V. et al. Flow of cortical activity underlying a tactile decision in mice. Neuron 81, 179–194 (2014).

9. Chen, T.-W., Li, N., Daie, K. & Svoboda, K. A Map of Anticipatory Activity in Mouse Motor Cortex. Neuron 94, 866–879.e4 (2017).

10. Inagaki, H. K., Fontolan, L., Romani, S. & Svoboda, K. Discrete attractor dynamics underlies persistent activity in the frontal cortex. Nature 566, 212–217 (2019).

11. Guo, Z. V. et al. Maintenance of persistent activity in a frontal thalamocortical loop. Nature 545, 181–186 (2017).

12. Economo, M. N. et al. Distinct descending motor cortex pathways and their roles in movement. Nature 563, 79–84 (2018).

13. Gao, Z. et al. A cortico-cerebellar loop for motor planning. Nature 563, 113–116 (2018).

14. Basso, M. A. & May, P. J. Circuits for Action and Cognition: A View from the Superior Colliculus. Annu Rev Vis Sci 3, 197–226 (2017).

15. Wolf, A. B. et al. An integrative role for the superior colliculus in selecting targets for movements. J. Neurophysiol. 114, 2118–2131 (2015).

16. May, P. J. The mammalian superior colliculus: laminar structure and connections. Prog. Brain Res. 151, 321–378 (2006).

17. Wurtz, R. H. & Goldberg, M. E. Activity of superior colliculus in behaving monkey. 3. Cells discharging before eye movements. J. Neurophysiol. 35, 575–586 (1972).

18. Hikosaka, O. & Wurtz, R. H. Modification of saccadic eye movements by GABA-related substances. I. Effect of muscimol and bicuculline in monkey superior colliculus. J. Neurophysiol. 53, 266–291 (1985).

19. Dorris, M. C., Paré, M. & Munoz, D. P. Neuronal activity in monkey superior colliculus related to the initiation of saccadic eye movements. J. Neurosci. 17, 8566–8579 (1997).

20. Horwitz, G. D. & Newsome, W. T. Separate signals for target selection and movement specification in the superior colliculus. Science 284, 1158–1161 (1999).

21. Felsen, G. & Mainen, Z. F. Neural substrates of sensory-guided locomotor decisions in the rat superior colliculus. Neuron 60, 137–148 (2008).

22. Felsen, G. & Mainen, Z. F. Midbrain contributions to sensorimotor decision making. J. Neurophysiol. 108, 135–147 (2012).

23. Duan, C. A., Erlich, J. C. & Brody, C. D. Requirement of Prefrontal and Midbrain Regions for Rapid Executive Control of Behavior in the Rat. Neuron 86, 1491–1503 (2015).

24. Kamigaki, T. & Dan, Y. Delay activity of specific prefrontal interneuron subtypes modulates memory-guided behavior. Nat. Neurosci. 20, 854–863 (2017).

25. Goard, M. J., Pho, G. N., Woodson, J. & Sur, M. Distinct roles of visual, parietal, and frontal motor cortices in memory-guided sensorimotor decisions. Elife 5, (2016).

26. Churchland, M. M., Yu, B. M., Ryu, S. I., Santhanam, G. & Shenoy, K. V. Neural variability in premotor cortex provides a signature of motor preparation. J. Neurosci. 26, 3697–3712 (2006).

27. Stachniak, T. J., Ghosh, A. & Sternson, S. M. Chemogenetic synaptic silencing of neural circuits localizes a hypothalamus→ midbrain pathway for feeding behavior. Neuron 82, 797– 808 (2014).

28. Gandhi, N. J. & Katnani, H. A. Motor functions of the superior colliculus. Annu. Rev. Neurosci. 34, 205–231 (2011).

29. Krauzlis, R. J., Lovejoy, L. P. & Zénon, A. Superior colliculus and visual spatial attention. Annu. Rev. Neurosci. 36, 165–182 (2013).

30. Song, J.-H., Rafal, R. D. & McPeek, R. M. Deficits in reach target selection during inactivation of the midbrain superior colliculus. Proc. Natl. Acad. Sci. U. S. A. 108, E1433– 40 (2011).

31. Rossi, M. A. et al. A GABAergic nigrotectal pathway for coordination of drinking behavior. Nat. Neurosci. 19, 742–748 (2016).

32. Zingg, B. et al. AAV-Mediated Anterograde Transsynaptic Tagging: Mapping Corticocollicular Input-Defined Neural Pathways for Defense Behaviors. Neuron 93, 33–47 (2017).

33. Tervo, D. G. R. et al. A Designer AAV Variant Permits Efficient Retrograde Access to Projection Neurons. Neuron 92, 372–382 (2016).

34. Green, D. M., Swets, J. A. & Others. Signal detection theory and psychophysics. 1, (Wiley New York, 1966).

35. Li, N., Daie, K., Svoboda, K. & Druckmann, S. Robust neuronal dynamics in premotor cortex during motor planning. Nature 532, 459–464 (2016).

36. Leichnetz, G. R., Spencer, R. F., Hardy, S. G. & Astruc, J. The prefrontal corticotectal projection in the monkey; an anterograde and retrograde horseradish peroxidase study. Neuroscience 6, 1023–1041 (1981).

37. Komatsu, H. & Suzuki, H. Projections from the functional subdivisions of the frontal eye field to the superior colliculus in the monkey. Brain Res. 327, 324–327 (1985).

38. Raybourn, M. S. & Keller, E. L. Colliculoreticular organization in primate oculomotor system. J. Neurophysiol. 40, 861–878 (1977).

39. Huerta, M. F., Krubitzer, L. A. & Kaas, J. H. Frontal eye field as defined by intracortical microstimulation in squirrel monkeys, owl monkeys, and macaque monkeys: I. Subcortical connections. J. Comp. Neurol. 253, 415–439 (1986).

40. Schnyder, H., Reisine, H., Hepp, K. & Henn, V. Frontal eye field projection to the paramedian pontine reticular formation traced with wheat germ agglutinin in the monkey. Brain Res. 329, 151–160 (1985).

41. Hanes, D. P. & Wurtz, R. H. Interaction of the frontal eye field and superior colliculus for saccade generation. J. Neurophysiol. 85, 804–815 (2001).

42. Katz, L. N., Yates, J. L., Pillow, J. W. & Huk, A. C. Dissociated functional significance of decision-related activity in the primate dorsal stream. Nature 535, 285–288 (2016).

43. Lakshminarasimhan, K. J., Pouget, A., DeAngelis, G. C., Angelaki, D. E. & Pitkow, X. Inferring decoding strategies for multiple correlated neural populations. PLoS Comput. Biol. 14, e1006371 (2018).

44. Hanks, T. D. et al. Distinct relationships of parietal and prefrontal cortices to evidence accumulation. Nature 520, 220–223 (2015).

45. Yartsev, M. M., Hanks, T. D., Yoon, A. M. & Brody, C. D. Causal contribution and dynamical encoding in the striatum during evidence accumulation. Elife 7, (2018).

46. Piet, A. T., Erlich, J. C., Kopec, C. D. & Brody, C. D. Rat Prefrontal Cortex Inactivations during Decision Making Are Explained by Bistable Attractor Dynamics. Neural Comput. 29, 2861–2886 (2017).

## Methods References

47. Zhong, L. et al. Causal contributions of parietal cortex to perceptual decision-making during stimulus categorization. Nat. Neurosci. 22, 963–973 (2019).

48. Guo, Z. V. et al. Procedures for behavioral experiments in head-fixed mice. PLoS One 9, e88678 (2014).

49. Appell, P. P. & Behan, M. Sources of subcortical GABAergic projections to the superior colliculus in the cat. J. Comp. Neurol. 302, 143–158 (1990).

50. Guizar-Sicairos, M., Thurman, S. T. & Fienup, J. R. Efficient subpixel image registration algorithms. Opt. Lett. 33, 156–158 (2008).

51. Harvey, C. D., Coen, P. & Tank, D. W. Choice-specific sequences in parietal cortex during a virtual-navigation decision task. Nature 484, 62–68 (2012).

